# Reprogramming of neuronal genome function and phenotype by astrocytes

**DOI:** 10.64898/2026.03.07.710282

**Authors:** Boxun Li, Kevin T. Hagy, Alexias Safi, Michael A. Beer, Alejandro Barrera, Sara Geraghty, Ruhi Rai, Alyssa N. Pederson, Samuel J. Reisman, Michael I. Love, Patrick F. Sullivan, Cagla Eroglu, Gregory E. Crawford, Charles A. Gersbach

## Abstract

Heterotypic cell-cell interactions are critical to governing cellular physiology, disease progression, and responses to the environment and pharmacologic interventions. For example, neurons and astrocytes engage in intricate interactions that are essential for brain development and function^1–3^. However, the transformation of these extracellular signals into epigenomic regulation that governs cell function is poorly understood. Here, we report that weeks of co-culture between human induced pluripotent stem cell (hiPSC)-derived neurons and mouse cortical astrocytes extensively reprograms gene expression and the chromatin accessibility landscape in neurons, affecting thousands of genes and putative gene regulatory elements (REs), including many transcription factors (TFs). These genes are enriched for functions implicated in neuronal differentiation and maturation, and tend to be impacted in schizophrenia, and autosomal dominant Alzheimer’s disease. Through complementary CRISPR interference and activation screens, we recapitulated hundreds of astrocyte-induced transcriptional and chromatin remodeling events in mono-cultured neurons at both promoters and distal regulatory elements (REs) of TF genes. We discovered functional REs for ∼50 astrocyte-responsive TF genes, providing a map of gene regulatory network control. Astrocyte-responsive TF genes fall into groups that exert independent or counter-balancing transcriptional effects, highlighting the complex coordination of the neuronal response to astrocytes. Functional effects of specific TFs, including POU3F2 and TFAP2E, on neurite morphology and neuronal electrophysiology are consistent with transcriptional effects, demonstrating the capacity of direct epigenetic control to mimic heterotypic cellular signals. This work illuminates the regulation of neurodevelopment-and disease-relevant gene modules by neuron-astrocyte interactions, and provides a blueprint for applying modern functional genomics to uncover the links between cell microenvironment and epigenomic programming.

**Highlights:**

- Neuronal gene expression and chromatin accessibility landscape are profoundly remodeled by astrocytes over weeks of co-culture
- Astrocyte-responsive neuronal gene modules and neuron-responsive astrocytic gene modules are enriched for genes associated with schizophrenia and familial Alzheimer’s Disease
- Single-cell CRISPR interference and activation screens of astrocyte-responsive gene regulatory elements identified dozens of functional regulatory elements of TF genes in neurons
- Single-cell CRISPR interference and activation screens of >200 astrocyte-responsive TF genes uncovered discrete functional clusters that promote neuronal maturity or stemness
- Astrocyte-responsive TF genes reprogram neuronal electrophysiology and neurite morphology

## Introduction

Since the completion of the Human Genome Project, over two decades of genomics research has transformed our understanding of genome structure and regulation^5^, leading to the first examples of the promise of genomic medicine in mitigating disease-causing genetic variants^6^. However, despite these advances and efforts of large consortia^7–9^, functional interpretation of the genome remains a major challenge. One key hurdle to a complete understanding of genome function is deciphering the context-dependent nature of gene expression regulation, including the impact of cell type, developmental stage, and microenvironment^10–12^.

Cell-cell interactions comprise a key component of the microenvironment in which diverse cell types interact within tissues. Neuron-astrocyte interactions (**NAI**s) exemplify the importance of cell-cell interactions in health and disease. Astrocytes are the most abundant glial cell type in the brain. Once thought to play a passive and supportive role for neurons, they have been shown to actively govern important neuronal processes such as the generation and function of synapses and regulation of metabolism that are essential to healthy neuronal function^1–3^. NAIs are disrupted in heritable neurodevelopmental and neurodegenerative diseases (e.g., Fragile X syndrome and Alzheimer’s disease) and are postulated to play a role in pathological progression^2,13,14^. However, the genomic reprogramming mediated by NAIs to impact cellular functions remains largely unmapped.

Here, we show that NAIs reprogram the epigenome, gene regulatory networks, and cellular functions of neurons through epigenetic and transcriptional events. Specifically, about a quarter of expressed genes (∼5000 of ∼22,000, FDR<0.01) and ∼10% of accessible chromatin regions (∼26,000 of ∼260,000, FDR<0.01) in neurons are responsive to astrocytes. Many of these astrocyte-responsive genes are disease-relevant. Through comprehensive CRISPR epigenome perturbations of a group of ∼200 astrocyte-responsive transcription factor (TF) genes, we recapitulated transcriptional events downstream of astrocyte signals and revealed how they mediate the broad changes in gene expression and chromatin accessibility downstream of NAIs. We further demonstrated that the astrocyte-responsive TF genes are in turn regulated by astrocyte-responsive distal REs. These epigenetic and transcriptional events form discrete groups that exert opposing effects while collectively improving neuronal health and maturity.

These discoveries highlight the importance of NAIs in the regulation of neurodevelopment- and disease-relevant gene modules, and provide a blueprint for applying CRISPR perturbation screens with single-cell transcriptome-wide readouts to uncover the links between cell environment and TF genes, REs, and cellular functions. These findings serve as a foundation for understanding how genome function, as well as the effects of genetic variation, can propagate from one cell type to another through heterotypic cell-cell interactions.

## Results

### NAIs extensively reprogram gene expression modules in neurons and astrocytes

#### NAIs reprogram gene expression in neurons

To model NAIs and study their genomic effects, we utilized an *in vitro* co-culture model of hiPSC-derived glutamatergic neurons induced by *Ngn2* overexpression and primary cortical astrocytes isolated from neonatal (∼postnatal day 1) mice^15–17^. Co-culture of human neurons and murine astrocytes have been widely used to model NAIs and to increase neuronal maturity^18–22^. The *Ngn2*-induced differentiation of iPSCs rapidly produces relatively homogenous cortical glutamatergic neurons^15,23^, matching the cortical origin of astrocytes. The neuronal differentiation goes through phases of pre-differentiation (mitotic, progenitor-like) and maturation (post-mitotic, neuronal)^16^. One day after entering the post-mitotic maturation phase, neurons were co-cultured with astrocytes at a ratio of five neurons to one astrocyte. We estimate this system to roughly model midgestation, when neurons first come into contact with astrocytes, based on the facts that the neurons are post-mitotic and initiating neurite outgrowth, and the approximate match between mouse neonatal and human midgestational brain development^24–26^. This model was chosen primarily based on common use, accessibility and tractability for complex genomic studies. As controls, the two cell types were cultured in isolation with the same seeding densities as in co-culture (**Fig. 1A**).

**Figure 1:**
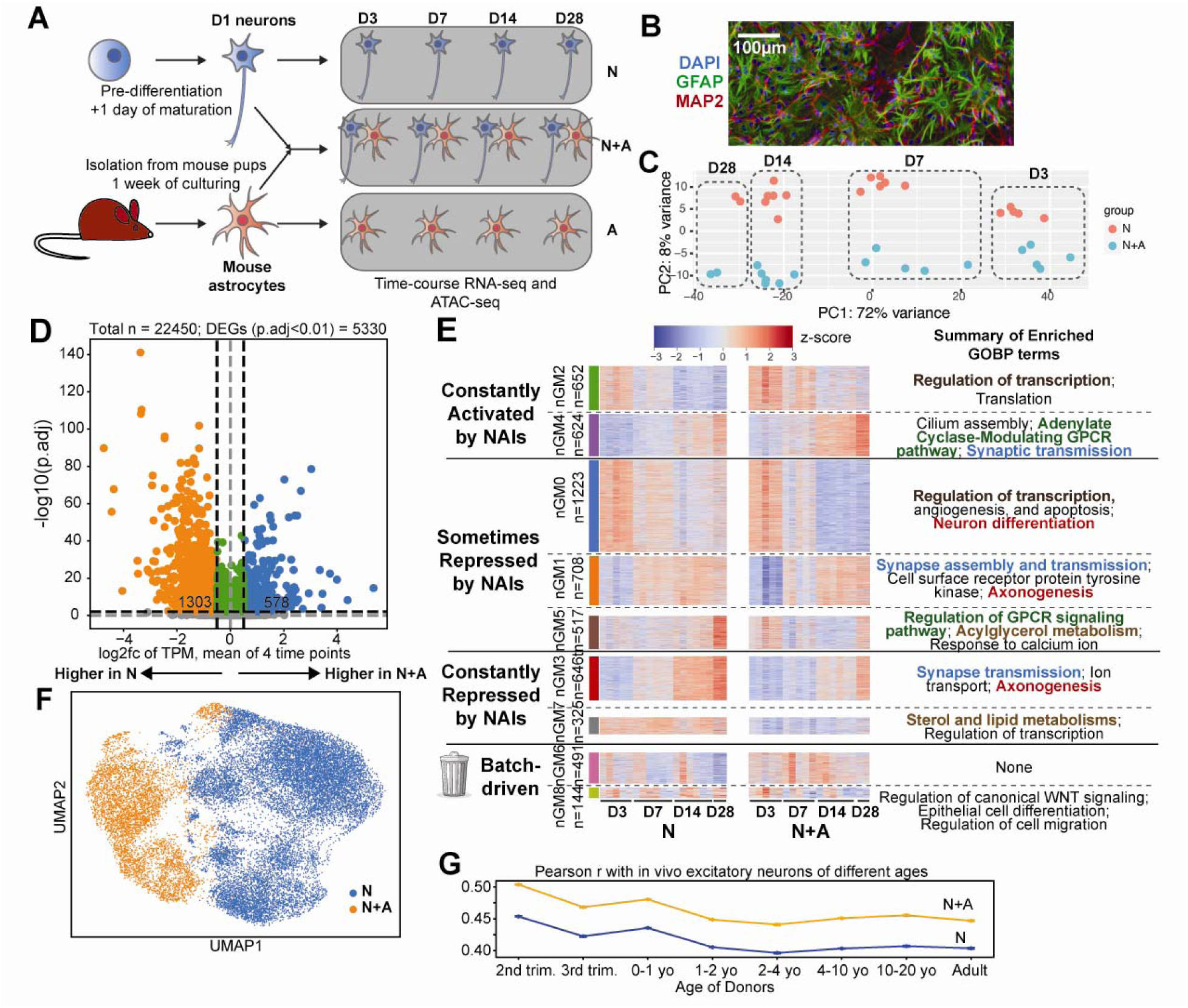
Neuron-astrocyte interactions reprogram neuronal gene expression programs. (A) Graphical illustration of the in vitro neuron-astrocyte co- and mono-cultures used to model and profile molecular phenotypes downstream of neuron-astrocyte interactions. (B) Immunofluorescence image of neuron-astrocyte co-culture on D7 showing DAPI, GFAP protein staining (astrocyte marker) and MAP protein staining (neuron marker). (C) PCA plot showing neuronal transcriptomes (PC1 vs PC2), marked by time points and colored by mono-(**N**) or co-culture (**N+A**) conditions. (D) Volcano plot showing the changes of astrocyte-responsive neuronal genes between N and N+A conditions, average of 4 time points. (E) Heatmap showing z-scaled expression of astrocyte-responsive neuronal genes across mono- and co-culture conditions, time points, and replicates, with enriched GOBP terms manually summarized. GOBP term colors: red, neuron differentiation-related; blue, synapse-related; green, GPCR-related; brown: metabolism-related. (F) UMAP plot showing mono-(**N**) or co-cultured (**N+A**) neurons profiled by scRNA-seq. (G) Line chart showing the mono-cultured (**N**) and co-cultured (**N+A**) neurons’ Pearson r with *in vivo* excitatory neurons of various donor ages from Velmeshev et al., 2023 using data shown in panel **E**.

We used bulk RNA-seq to investigate the impact of NAIs on gene expression in mono-cultured neurons (**N**), co-cultured neurons and astrocytes (**N+A**), and mono-cultured astrocytes (**A**) on days 3, 7, 14, and 28 after entering maturation (i.e., D3, D7, D14, and D28) (**Fig. 1A, B, Table S1**). Because of the ∼80 million-year mouse-human evolutionary distance, RNA-seq reads can be assigned to an astrocyte (mouse alignment) or neuron (human alignment) with high confidence (>99.9%)^27^. We then tested for differential gene expression between neurons in **N** vs **N+A** (only analyzing human reads) and astrocytes (**A** vs **N+A**, only analyzing mouse reads)^27–29^ (**Extended Data Fig. 1A**).

To investigate the neuronal transcriptional response to astrocytes, we performed principal component analysis (PCA). Most variation in neuronal transcriptomes was attributed to time point and co-culture condition (**Fig. 1C**). From D3 to D28, 5330 genes (23.7% or 5330/22450, FDR < 0.01) were differentially expressed between N and N+A (**Fig. 1D, Table S2, Methods**). We termed these genes astrocyte-responsive neuronal (**ArN**) genes. Notably, the responses of ArN genes are highly correlated in consecutive time points (mean Pearson r = 0.63), but uncorrelated between the first and last time points (Pearson r = −0.02), highlighting their temporal dynamics and the need for capturing multiple time points (**Extended Data Fig. 1B**).

Since ArN gene expression changes over time, we aimed to group genes into gene modules that reflect time- and co-culture condition-based expression patterns and biological processes. Using unsupervised clustering (**Methods**), we identified 9 gene modules that we termed **n**euronal **G**ene **M**odule 0 through 8 (**nGM**0-8), each representing a distinct expression pattern (**Fig. 1E**, **Extended Data Fig. 1C**; gene modules were ordered by size, with nGM0 being the biggest). For example, nGM2 represents genes that are consistently activated by astrocytes, while being silenced over time. By contrast, nGM7 is repressed by astrocytes, but stays constant over time. Over-representation analysis (**Methods**) (**Table S3**) revealed enrichment of Gene Ontology Biological Process (**GOBP**) terms related to neuron differentiation and axonogenesis (nGM0, 1, 3), synaptogenesis and synapse transmission (nGM1, 3, 4), GPCR signaling pathways (nGM4, 5), and lipid metabolism (nGM5, 7), which are all pathways known to be influenced by NAIs ^30^. We also found enrichment for transcriptional regulation processes, including many TF and epigenetic modifier genes, in nGM0 and 2, suggesting they may act as drivers of the broad transcriptional reprogramming.

#### NAIs reprogram gene expression in astrocytes

We next investigated how astrocytes respond to neurons. To confirm the purity of astrocytes isolated from mouse cortices, we found the median expression of six astrocyte marker genes to be >10-fold relative to markers of any potential contaminating cell types (**Extended Data Fig. 2A**, **Table S4**). We found that NAIs also extensively rewire the gene expression programs of astrocytes, influencing 3,587 genes over four weeks, which we termed neuron-responsive astrocytic (**NrA**) genes (**Extended Data Fig. 2B**; **Table S5**). These include well-studied astrocyte genes that are involved in recycling of the excitatory neurotransmitter glutamate (e.g., *Slc1a2*) and regulation of synaptogenesis (e.g., *Thbs*)^30,31^. However, unlike ArN genes in neurons, the responses of NrA genes are highly correlated between both consecutive (mean Pearson ⍴ = 0.78) and the first and last time points (⍴ = 0.51) (**Extended Data Fig. 2C**), suggesting modest temporal dynamics. Consistent with this, unsupervised clustering of NrA genes into 7 **a**strocytic **G**ene **M**odules (**aGM**0-6) revealed less complex expression patterns than those found in neurons, with 6 of 7 aGMs either consistently activated or repressed across all time points (**Extended Data Fig. 2D**). GOBP terms, including nervous system development, synaptic transmission, and cholesterol metabolism, are enriched in different aGMs (**Table S6**). These are consistent with known functions of astrocytes, and overlap those of the nGMs, suggesting coordinated regulation of shared biological processes in both cell types.

#### Neuronal gene expression reprogramming is confirmed by single-cell RNA-seq

To corroborate the findings above, we performed single-cell RNA-seq (scRNA-seq) on mono-and co-cultured neurons (**N** and **N+A**) at D14 using a droplet-based method (10X Genomics 5’ HT v2). We could distinguish between droplets that contained only neurons, only astrocytes, or both by the proportions of reads mapped to the human and mouse genomes (**Extended Data Fig. 3A, B**, **Methods**). After removing all astrocyte-containing droplets, comparison of droplets with only neurons in the N and N+A conditions identified 15 cell clusters (**Extended Data Fig. 3C**) and confirmed the previously reported cortical glutamatergic neuron fate (**Extended Data Fig. 3C, D, E**)^23^. Strikingly, mono- and co-cultured neurons clustered distinctly (**Fig. 1F**), mirroring the bulk RNA-seq results (**Fig. 1C**) and suggesting that neurons respond relatively homogeneously to astrocytes. Differential expression analysis identified 6,921 differentially expressed genes (DEGs; FDR < 0.01; **Methods**) between mono- and co-cultured neurons and their changes are highly correlated between single-cell and bulk (D14) measurements (Pearson r = 0.78, **Extended Data Fig. 3F**). We next compared the ArN genes with genes previously reported to be influenced transcriptionally in hiPSC-derived neurons by glial cells in two different settings^19,20^ and found significant overlaps (mean ∼1.5-fold enrichment; **Extended Data Fig. 1D**). A core set of 303 DEGS were identified in all three studies. These 303 genes were enriched for GOBP terms related to synaptic and nervous system development (**Extended Data Fig. 1E**, **Discussion**). Similarly, we found significant overlap of NrA genes with genes reported to be downstream of NAIs or involved in astrocytic maturation^32,33^ (∼2.6-fold enrichment **Extended Data Fig. 2E**).

#### NAIs increase neuronal transcriptional maturity

To test our hypothesis that astrocytes increase neuronal maturity, we correlated our neuron scRNA-seq data (shown in **Fig. 1F**) with excitatory neurons from a human cortical brain single-nucleus RNA-seq atlas containing both developmental and adult tissues, using Pearson correlation calculated from ∼17,000 genes that are common to both datasets (**Methods**)^34^. Both mono- and co-cultured neurons are most highly correlated with neurons from second trimester tissues, confirming their midgestational maturity (**Fig. 1G**). This is expected since iPSC-derived neurons are known to have fetal-like characteristics^35^. Moreover, we found that co-culture with astrocytes makes neurons more highly correlated with *in vivo* neurons (∼10% increase in Pearson r), and that co-cultured neurons correlate with *in vivo* adult neurons to a similar extent as mono-cultured neurons correlate with *in vivo* prenatal neurons. This suggests astrocytes provide general microenvironmental cues that make neurons transcriptionally more mature.

#### Response to NAIs prioritizes neuron-astrocyte communication signals

To understand the molecular interactions that could mediate NAIs, we inferred 83 pairs of potential ligand-receptor interactions between astrocytes and excitatory neurons based on gene expression data from the human brain single-nucleus RNA-seq dataset with CellChat^34,36^ (**Table S7**). We reasoned that for these putative ligand-receptor pairs, responsiveness to NAIs at the mRNA level (as highlighted in **Fig. 1D** and **Extended Data Fig. 2B**) could serve as functional evidence of a true interaction. We observed that 17 of 37 (46%), and 27 of 40 (68%), of the inferred interacting genes in neurons and astrocytes, respectively, are transcriptionally responsive to co-culture, supporting the roles of these 44 genes in NAIs (**Extended Data Fig. 2F**). Furthermore, we identified 21 interacting pairs where both the ligand and the receptor are responsive to NAIs in the respective cell type, including 3 astrocyte-neuron and 18 neuron-astrocyte ligand-receptor pairs (**Extended Data Fig. 2G**). Experimental evidence from the literature supports the physical interaction and disease relevance of some of these pairs, including astrocyte:PTN-neuron:PTPRZ interaction in major depressive disorder and neuron:NRG1-astrocyte:ERBB4 interaction in schizophrenia and Alzheimer’s Disease^37,38^. These analyses nominate known and potentially new molecular interactions that might mediate NAIs. Together, our results describe extensive reprogramming of gene expression programs that occur in both neurons and astrocytes when co-cultured. This phenomenon is broad and temporally dynamic, and results in more mature neurons that better mimic their *in vivo* counterparts.

### NAIs extensively rewire the neuronal chromatin landscape

Because the remodeling of the epigenome is central to gene regulation and plays important roles in brain development and disease^39^, we next investigated whether the NAI-induced reprogramming of gene expression modules is accompanied and reflected in the epigenome. We specifically focused on chromatin accessibility as a readout as it marks potential gene regulatory elements (RE). We performed bulk ATAC-seq analysis of mono- and co-cultured neurons and astrocytes over time (D3, D7, and D14), similar to that used for RNA-seq above (**Fig. 1A**, **Extended Data Fig. 4A, Table S1**, **Methods**). Sequence reads were similarly assigned to the human (neuron) or the mouse (astrocyte) genome to assign cell types, and data were of high quality for mono- and co-cultured neurons except for D28 samples, which were removed from downstream analysis beyond peak calling (**Extended Data Fig. 4F**, **Discussion**).

Most of the variability in neuronal chromatin accessibility can be attributed to time point and co-culture condition (**Fig. 2A**), indicating that chromatin accessibility is dramatically rewired by NAIs alongside gene expression. We detected a total of 26,429 **a**strocyte-**r**esponsive **n**euronal ATAC-seq **peaks** (**ArN peaks**) (FDR < 0.01) over the time course (**Fig. 2B, Table S8**), which accounted for ∼10% of all ATAC-seq peaks (n=265,420). Similar to RNA-seq, the changes of the ArN peaks are more highly correlated between consecutive time points (D3 vs D7 and D7 vs D14, mean Pearson r = 0.47) than between D3 and D14 (Pearson r = 0.15) (**Extended Data Fig. 4B**). These results indicate extensive and temporally dynamic rewiring of the neuronal epigenome in response to astrocytes.

**Figure 2:**
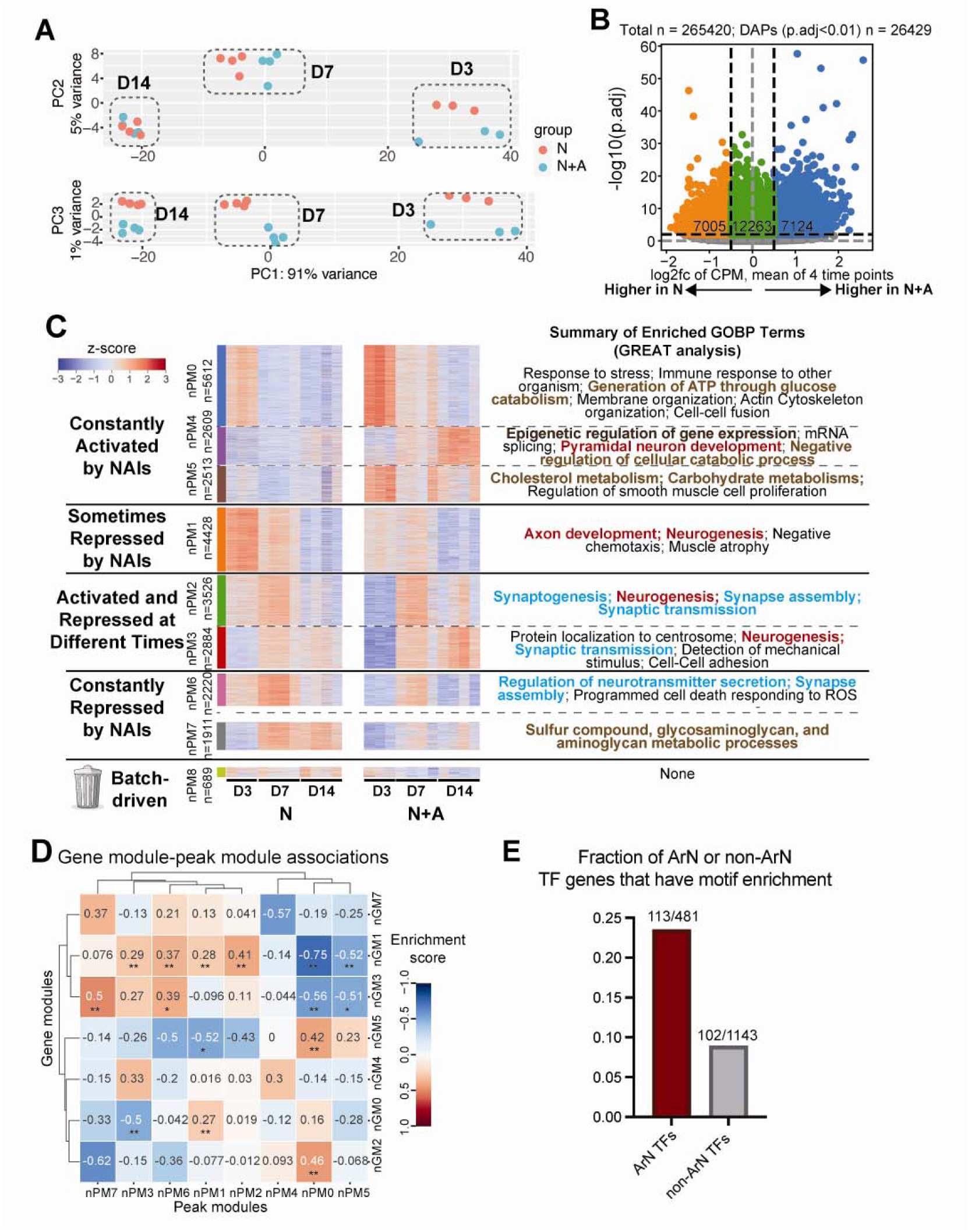
Neuron-astrocyte interactions reprogram neuronal epigenome. (A) PCA plot showing neuronal ATAC-seq profiles (top, PC1 vs PC2; bottom, PC1 vs PC3), marked by time points and colored by mono-(**N**) or co-culture (**N+A**) conditions. (B) Volcano plot showing the changes of astrocyte-responsive neuronal ATAC-seq peaks between N and N+A conditions, average of 3 time points. (C) Heatmap showing z-scaled accessibility of astrocyte-responsive neuronal ATAC-seq peaks across mono- and co-culture conditions, time points, and replicates, with enriched GOBP terms manually summarized. GOBP term colors: red, neuron differentiation-related; blue, synapse-related; green, GPCR-related; brown: metabolism-related. (D) Heatmap showing the enrichment score of proximity-based associations between genes from a given nGM and peaks from a given nPM. Enrichment score is defined as log2(observed/expected) total counts of peaks in the given nPM for genes in the given nGM. *: p<0.05; **: p<0.001. (E) Barplot showing the fraction of ArN and non-ArN TF genes that have motif enrichment in ArN peaks.

We then investigated how the RNA-seq and ATAC-seq data correlated in the co-culture model. Unsupervised clustering of the ArN peaks (**Methods**) identified 9 clusters of ATAC peaks (termed **n**euronal **P**eak **M**odule 0-8, or nPM0-8) (**Fig. 2C**, **Extended Data Fig. 4C**). nPMs identify groups of ATAC peaks that are similarly impacted by astrocyte co-culture at different time points. By performing Genomic Regions Enrichment of Annotations Tool (GREAT) ^40,41^ analysis on nPMs (**Methods, Table S9**), we identified enriched GOBP terms that were similarly detected from RNA-seq data, including those related to neurogenesis and synaptogenesis (nPM1, 2, 3, 6), synaptogenesis and transmission (nPM 2, 3, 6), metabolism (nPC0, 4, 5, 7), and transcriptional regulation (nPC4). By associating each ATAC-seq peak to its nearest gene, we found that ArN genes repressed in co-culture are on average more likely to be near ArN ATAC-seq peaks that lose accessibility in co-culture, while ArN genes that are activated are more likely to be near ArN ATAC-seq peaks that gain accessibility (**Methods**, **Extended Data Fig. 4D**). Furthermore, to uncover higher-resolution gene regulatory relationships between chromatin landscape and gene expression, we tested for the associations between all possible nGM-nPM pairs (n=56 pairs, 7 nGMs x 8 nPMs) using the same nearest gene-based approach. 15 associations were found to be statistically significant (**Fig. 2D**) (**Methods**). As one example, the nGM2 module represents genes that are consistently activated by astrocytes across the time course, and has significant positive association with chromatin accessibility module nPM0, which gains chromatin accessibility in the presence of astrocytes. Thus, nGM2 and nPM0 exhibit concordant responses to astrocytes. Taken together, these results support that ArN ATAC peaks likely represent REs of ArN genes.

To mechanistically understand the association between gene expression and chromatin accessibility, we used gkm-SVM to nominate transcription factors (TFs) in each nPM that likely drive chromatin accessibility changes^42,43^ (**Extended Data Fig. 4E**, **Table S10**). For example, the NEUROD1 motif was the second most important motif for the nGM2-nPM0 association. The NEUROD1 motif was predicted to be bound by four astrocyte-responsive TFs (NEUROD1, NEUROD4, NEUROG1, and OLIG3) that also belong to nGM2, providing a potential mechanism for the self-regulation of nGM2. Overall, we observed that expressed TFs that are astrocyte-responsive at the mRNA level are more likely than unresponsive ones to be nominated as regulators of chromatin accessibility modules (2.6-fold enrichment) (**Fig. 2E**), supporting that astrocytes rewire the neuronal epigenome through regulating TF gene expression.

In summary, in addition to gene expression programs, astrocytes also reprogram the neuronal epigenome, and these two phenomena are concordant at both individual gene/element and module levels. Furthermore, our data suggest that these phenomena are jointly mediated by astrocyte-responsive TFs (**Extended Data Fig. 5A**), positioning them as candidates for the further investigation of the gene regulatory mechanisms underlying the reprogramming of neuronal epigenome and transcription.

### Perturbation of promoters and putative REs of TFs decodes the gene regulatory logic governing neuronal response to astrocytes

To identify protein-coding and non-coding drivers of the molecular phenomena downstream of NAIs, we focused on the 222 TF genes expressed by neurons that are strongly responsive (|log_2_fc|>1) to astrocytes at the mRNA level at one or more time points (**Extended Data Fig. 5A, Table S11**) and their distal REs (≤100kb from TSS). We used high-throughput CRISPR interference and activation (CRISPRi/a) epigenome editing paired with scRNA-seq (i.e., Perturb-seq) to interrogate the functions of the promoters and distal REs of these TF genes (**Fig. 3A**, **Fig. 4A**). In total, we identified 280 promoter targets and 735 non-promoter astrocyte-responsive neuronal ATAC-seq peaks as putative RE (pRE) targets (**Extended Data Fig. 7A, Extended Data Fig. 5B**; **Methods**).

**Figure 3:**
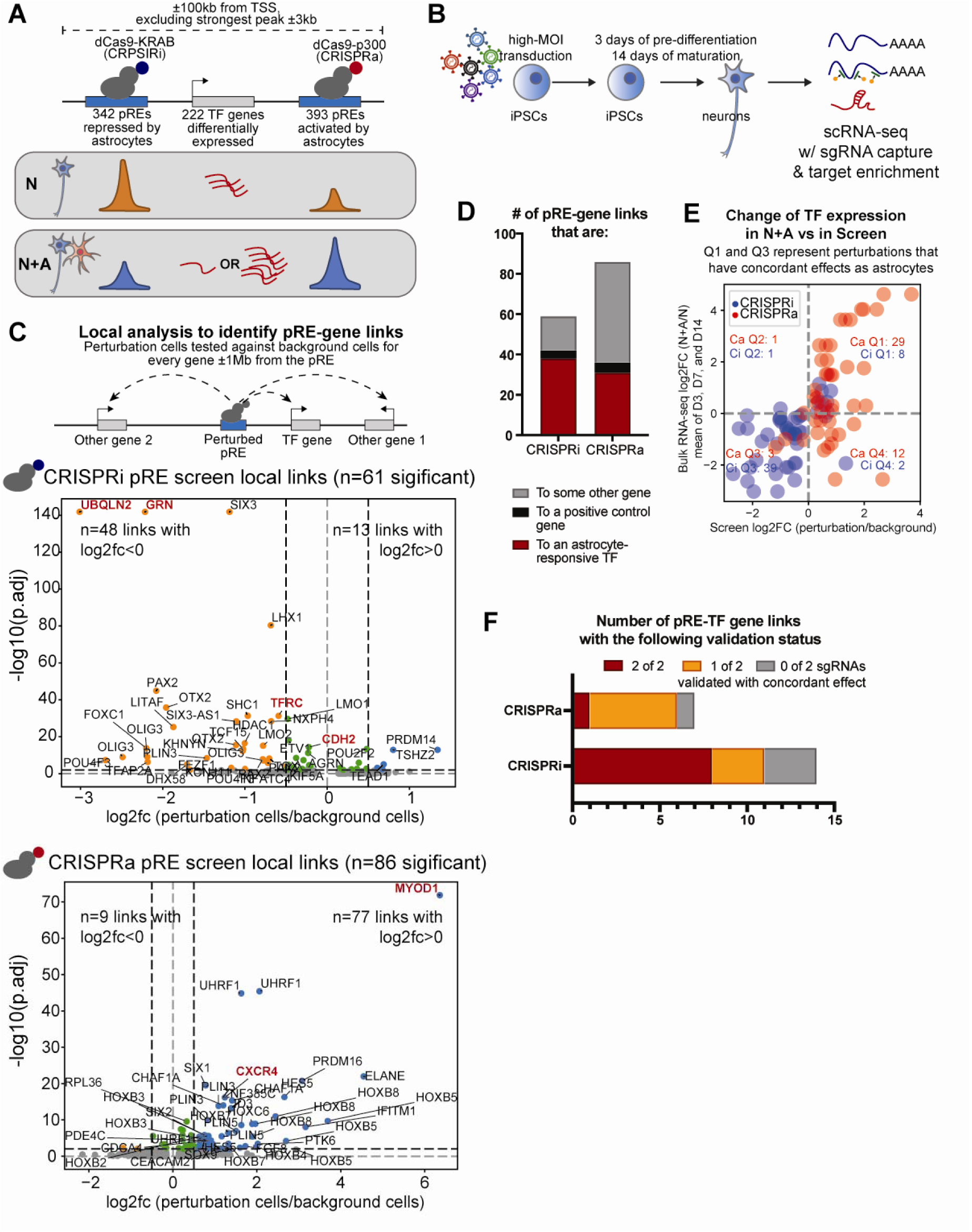
Single-cell pRE Perturb-Seq reveals the cis regulatory landscapes of ∼50 astrocyte-responsive TFs. (A) Schematics showing the design of the CRISPRi/a (by dCas9-KRAB and dCas9-p300, respectively) Perturb-seq screens of 735 ArN pREs that are near n=222 ArN TF genes. 342 pREs become more closed with NAIs, which are targeted by CRISPRi (KRAB); 393 pREs become more open with NAIs, which are targeted by CRISPRa (p300). (B) Schematics showing the pRE Perturb-seq experimental setup. (C) Schematics representing local analysis (top), and volcano plots (middle and bottom) summarizing all local fRE-gene links that were discovered. Middle, fRE-gene links from the CRISPRi pRE screen. Bottom, fRE-gene links from the CRISPRa pRE screen. Red and bolded labels indicate positive controls. (D) Stacked barplot showing the counts of local pRE-gene links to an ArN TF gene (those with FDR<0.01 **Extended Data** Fig. 5A), a positive control gene, or a gene not of these two categories. (E) Scatterplot showing the direction of change of genes with fRE CRISPRi or CRISPRa perturbation, vs with NAIs. (F) Barplot showing bulk RT-qPCR validation results of individual sgRNAs selected for 14 and 7 fREs-gene pairs for CRISPRi and CRISPRa screens, respectively.

**Figure 4:**
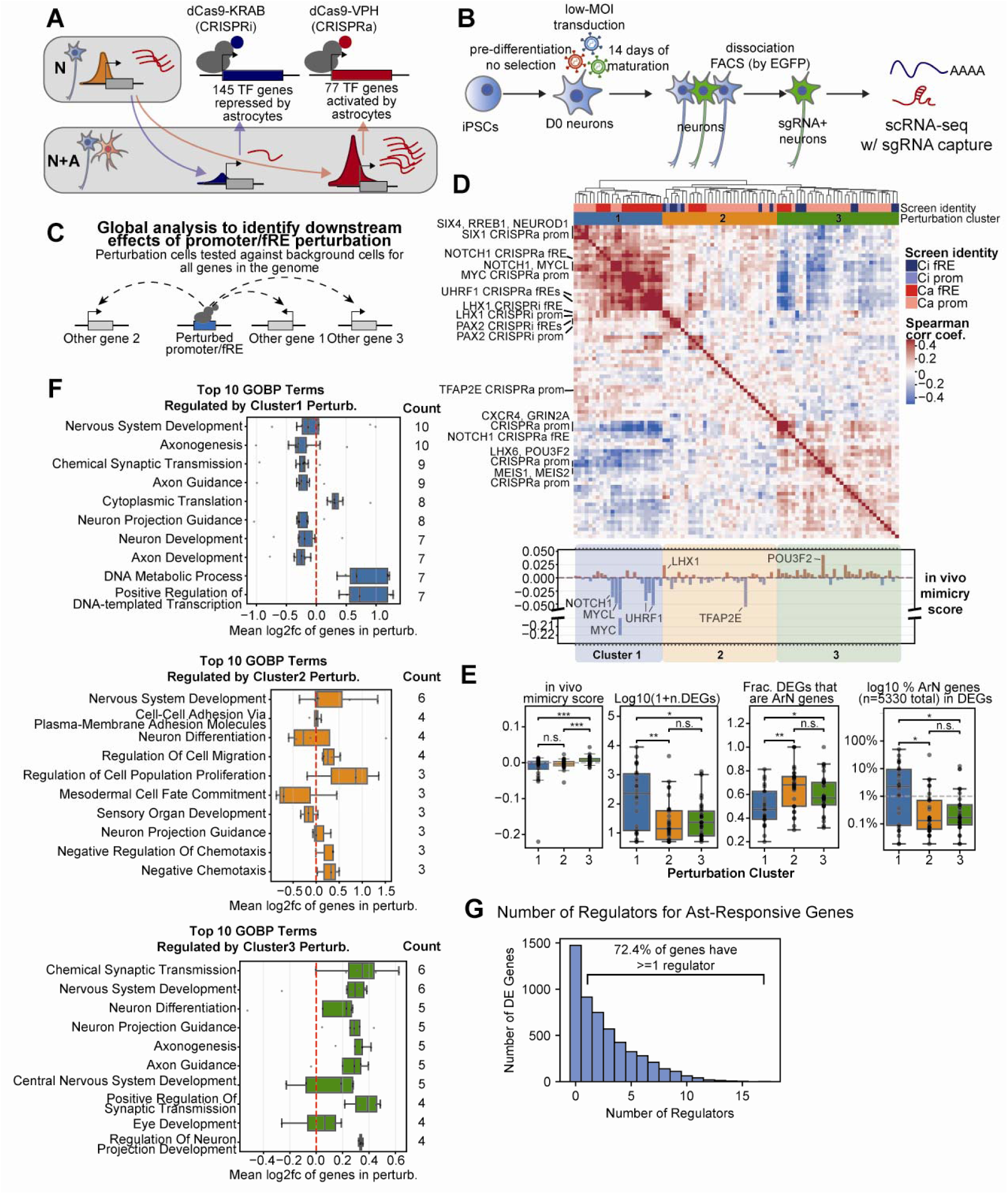
Single-cell promoter Perturb-Seq reveals counter-balancing effects of discrete groups of perturbations. (A) Schematics showing the design of the CRISPRi/a (by dCas9-KRAB and dCas9-VPH, respectively) Perturb-seq screens of ArN TF gene promoters. (B) Schematics showing the promoter Perturb-seq experimental setup. (C) Schematics representing global analysis. (D) **Top**, heatmap showing the pairwise Spearman correlation of 88 perturbations from the 4 Perturb-seq experiments, with columns color-coded by screen identity (first row) and perturbation cluster (second row), and with example perturbations pointed out. Perturbations fall into four clusters (1-3). **Bottom**, barplot showing perturbations in the same order as the columns of the heatmap, showing in vivo mimicry score of the individual perturbations. (E) Box plots showing the distributions across the three perturbation clusters of a) in vivo mimicry score, b) log10(number of global DEGs + 1), c) Fraction of global DEGs that are ArN genes, and d) % of ArN genes covered by the global DEGs of this perturbation across the four perturbation clusters. Mann-Whitney U test results are reported (based on raw p values) between perturbation clusters: n.s.: p>0.05; *: p<0.05; **: p<0.01; ***: p<0.001. The individual perturbation-level description of these characteristics is shown in **Extended Data** Fig. 8A. (F) Boxplots showing the mean log2FC of genes belonging to the top 10 GOBP terms most frequently regulated by perturbations in cluster 1 (top, blue), 2 (middle, orange), and 3 (bottom, green), respectively. On the right of the boxplots are shown the counts of members within each perturbation cluster whose downstream genes are significantly enriched for that GOBP term. (G) Histogram showing the number of regulators per astrocyte-responsive gene found among perturbations shown in Fig. 4D.

To mimic the transcriptional and epigenetic events elicited by astrocytes, we perturbed mono-cultured neurons using CRISPRi on TF promoters and pREs that are repressed by astrocytes, and CRISPRa on TF promoters and pREs that are activated by astrocytes. This approach allows us to gain causal insights into *cis* and *trans* gene regulatory logic of astrocyte-responsive TFs, expand the catalog of experimentally established functional REs in human neurons, and nominate TF genes for epigenetic engineering that may synthetically recapitulate the effects of astrocytes. Below, the results of these experiments are shown in the order of 1) *cis* regulatory logic and 2) *trans* regulatory logic.

### Single-cell analysis of pRE perturbations reveals the cis regulatory landscapes of ∼50 astrocyte-responsive TFs

To chart the cis regulatory landscape of ArN TFs, we performed CRISPRi/a Perturb-seq in mono-cultured neurons to interrogate all ArN pREs (i.e., ATAC-seq peaks, **Fig. 2B**) located ≤100kb from any TSS of the 222 ArN TF genes, excluding the strongest promoter for each gene as promoters were addressed in separate experiments, for a total of 735 targets (**Fig. 3A**, **Extended Data Fig. 5A**, **Methods**). Of these, 342 and 393 were targeted by CRISPRi (dCas9-KRAB) and CRISPRa (dCas9-p300), respectively (**Extended Data Fig. 5B**). We found that astrocyte-activated TF genes tend to have more local ATAC-seq peaks that become more accessible (CRISPRa pRE targets), and conversely, astrocyte-repressed TFs tend to have more local peaks that become less accessible (CRISPRi pRE targets) (**Extended Data Fig. 5C**), consistent with the overall correlation of effect direction of ArN genes and peaks (**Extended Data Fig. 4D**). For CRISPRi, we used a WTC-11 iPSC line that stably expresses dCas9-KRAB^16^. For CRISPRa, we engineered a WTC-11 iPSC line to stably express dCas9-p300, and verified that it activates gene expression in differentiated neurons at both promoter and distal regulatory regions^4,44^ (**Extended Data Fig. 5D**).

We designed ∼10 sgRNAs/target for each of the 735 pRE targets, along with positive and negative control sgRNAs (**Fig. 3B**, **Extended Data Fig. 5B**, **Methods**). Notably, we applied target enrichment of the 222 TF genes of interest to increase statistical power of downstream analysis^45^. The enriched and unenriched transcriptome libraries were sequenced and combined for data analysis (**Extended Data Fig. 5H**). After preprocessing and quality filtering, we retained 13,382 and 17,887 high-quality cells that have at least 1 sgRNA assignment in the CRISPRi and CRISPRa screens, respectively. We observe a median of 5 and 3 sgRNAs/cell, and a median of 1668 and 966 cells per target region, respectively (**Extended Data Fig. 5B**).

We identified 57 and 55 unique target regions as functional gene regulatory elements in *cis* (distance ≤1 Mb from the perturbed site, FDR < 0.01, effect size > 10%), which we termed functional REs (fREs) (**Fig. 3C**, **Table S12**), using established methods^46,47^. As expected, most CRISPRi perturbations resulted in down regulation and most CRISPRa perturbations resulted in up regulation. Importantly, these discoveries include 4 of 4 and 4 of 5 positive controls, respectively, in the CRISPRi screen (promoters of *UBQLN2*, *GRN*, *TFRC*, *CDH2*) and CRISPRa screen (promoters of *MYOD1*, *CXCR4*, *GRIN2A* and *COL4A1*) (**Fig. 3C**, red, bolded labels), indicating robustness of the results.

Besides the positive controls, the CRISPRi screen linked 53 fREs to 45 unique genes (ArN TF genes as well as other genes; same below) for a total of 57 fRE-gene links, with a mean fRE-to-gene distance of 26kb. The CRISPRa screen linked 51 fREs to 62 unique genes for total of 81 fRE-gene links, with a mean fRE-to-gene distance of 83kb (**Extended Data Fig. 5E**). Most fREs regulated only one gene, and most genes were regulated by only one fRE. However, the *cis* regulatory logic is very complex in some cases (**Extended Data Fig. 5F**). As examples, the CRISPRi screen discovered 3 fREs for the TF gene SIX3, two of which also regulate the antisense transcript SIX3-AS1 (**Extended Data Fig. 6A**). Similarly, in the CRISPRa screen, 7 fREs were discovered in the HOXB locus, with HOXB2, HOXB3, HOXB5, HOXB7, HOXB8, and HOXB-AS1 each regulated by at least two of them (**Extended Data Fig. 6B**). Substantial proportions of the fRE-gene links (67% in CRISPRi screen and 38% in CRISPRa screen) are linked to ArN TF genes (**Fig. 3D**). In total, at least one fRE was discovered for 49 of the 222 ArN TFs (22.1%). Moreover, of the links from the CRISPRi and CRISPRa screens that involve an ArN gene (shown in **Fig. 1D**), 94% and 71% showed concordant effects with the influence of astrocytes (**Fig. 3E**), highlighting that our CRISPRi/a approach provides a reliable framework to recapitulate the epigenetic mechanisms underlying the neuronal response to astrocytes.

To validate the functions of fREs, we sampled 14 CRISPRi and 7 CRISPRa fRE-TF gene links across a range of statistical significance. For each fRE, we selected two sgRNAs and tested them individually in mono-cultured neurons by RT-qPCR (**Methods**). When compared to a non-targeting sgRNA control, 11 of 14 (79%) and 6 of 7 (86%) fREs have at least one sgRNA that validates with concordant effect with the single-cell analysis (**Fig. 3F**, **Table S13**), confirming the validity of the discoveries. Notably, one fRE CRISPRi perturbation upregulated PRDM14 in both RT-qPCR and Perturb-seq (**Fig. 3C, top**), indicating that the perturbations that showed unexpected direction of effect could be biologically meaningful. It is also noteworthy that perturbations of different fREs of the same TF gene often led to different levels of regulation, providing ways to tune TF expression (**Extended Data Fig. 5G**).

To evaluate disease relevance, we queried the fRE-linked genes against the ClinGen database, and found that 12 and 8 fRE-linked genes from the CRISPRi and CRISPRa pRE screens respectively, excluding positive controls, are associated with disease (**Table S14**), suggesting that the fREs could be exploited in the future to engineer the expression these genes for therapeutic purposes.

In summary, by using complementary CRISPRi/a screens of 735 pREs, we discovered 104 fREs, including ones that regulate 49 ArN TF genes. Epigenome editing robustly recapitulated the epigenetic events downstream of NAIs, and revealed the sometimes complex *cis* gene regulatory logic of the neuronal response to astrocytes. Our approach provides a blueprint for studying *cis* gene regulation in cell-cell interactions, and creates future opportunities to epigenetically engineer disease-relevant TFs.

### Promoter perturbations reveal counter-balancing effects of discrete groups of TFs

To map the downstream gene programs regulated by each of the ArN TF genes, we targeted the promoter region(s) of every TF with Perturb-seq CRISPRi (dCas9-KRAB) or CRISPRa (dCas9-4xVP48-P65-HSF1 (dCas9-VPH)) using previously reported cell lines (**Fig. 4A**, **Extended Data Fig. 5A, Extended Data Fig. 7A, B**)^16,48^. We targeted 10 gRNAs to each promoter region. We transduced WTC11 neural progenitor-like cells with lentiviral sgRNA libraries at low MOI, differentiated them into neurons, sorted neurons that expressed the lentiviral construct by fluorescence-activated cell sorting (FACS), and performed scRNA-seq (**Fig. 4B**). We retained 67,176 and 80,539 high-quality cells with a median of 1 sgRNA/cell and a median of 474 and 1155 cells per promoter for the CRISPRi and CRISPRa screens, respectively (**Extended Data Fig. 7B**). Strikingly, in the CRISPRa screen, cells with a MYC promoter sgRNA have ∼10x as many cells compared to other sgRNAs, consistent with MYC being a proto-oncogene and a positive regulator of proliferation^49^ (**Extended Data Fig. 7C**). To avoid statistical bias, we downsampled the cells containing a MYC sgRNA to match the other perturbations (**Methods**). For each perturbation, we characterized the downstream genome-wide transcriptional profiles (**Fig. 4C**, **Extended Data Fig. 5H**, **Table S15**).

To verify robustness of the screen, we found that CRISPRa resulted in significant (*p* < 0.05) upregulation of 71 of the 76 targeted TF genes (93%), as well as all 4 positive controls (**Extended Data Fig. 7D**). CRISPRi, however, only significantly down-regulated 50 of the 122 (41%) targeted TF genes that were tested and only 2 of 4 positive control genes (UBQLN2 and GRN were significant; TFRC and CDH2 were not). Notably, the successfully repressed targets have ∼4.4x the expression levels of the unsuccessful targets at baseline (**Extended Data Fig. 7E**, median CPM 7.9 vs 1.8), suggesting that low statistical power played a role for the difficulty in detecting the repression of more lowly expressed genes.

To define global downstream regulatory networks, we focused on CRISPRi/a promoter perturbations that significantly down- or up-regulated the intended target gene, respectively, as well as fRE perturbations (i.e., pREs linked to a local target gene), because they have a clear mechanism of downstream gene regulatory effects. We further restricted downstream analysis to only those perturbations that led to more than 3 differentially expressed genes (DEGs; FDR < 0.01). By these criteria, 88 perturbations across the 4 screens were included, including 10 positive controls. In total, we discovered 36,480 global perturbation-gene links, 1,617 of which are from positive controls (**Extended Data Fig. 7F**). There is a strong disparity between CRISPRi and CRISPRa, with the two CRISPRa screens accounting for the vast majority of total discoveries. However, the number of total discoveries is driven by a handful of outliers, and the medians across the 4 screens are similarly moderate (range, 17 - 47.5 DEGs) (**Extended Data Fig. 7G**).

To understand the relationships between the TFs’ downstream gene regulatory networks, we performed pairwise correlation on the 88 perturbations across 247 genes that are DEGs of at least one perturbation in at least 3 of the 4 Perturb-seq datasets (**Methods**). This selection step minimizes the chance of including genes that were driven by experiment-specific batch effects. We found expected relationships, such as concordance between multiple perturbations of the same TF genes (e.g., CRISPRi perturbations of the fRE and promoter of LHX1, and CRISPRi perturbations of two different PAX2 fREs), as well as the known concordant functions between MYC and MYCL^49^. The perturbations fall into three clusters that exert opposing (cluster 1 vs 3) or largely independent (cluster 2 vs cluster1/3) effects (**Fig. 4D**, heatmap; **Table S16**). The perturbation clusters are distinguished by four metrics (**Fig. 4D** bar plot, **Fig. 4E**, and **Extended Data Fig. 8A**). Cluster 3 stands out with an overwhelmingly positive similarity score to *in vivo* excitatory neurons, or *in vivo* mimicry score, measured by correlation of the perturbed transcriptome to *in vivo* excitatory neurons relative to the background transcriptome from the same screen (**Methods**)^34^. This is consistent with the increased similarity of co-cultured neurons to *in vivo* neurons (**Fig. 1F**). The perturbation with the strongest *in vivo* mimicry score is the promoter activation of POU3F2 (also known as BRN2), a TF with well-established functions in brain development that inhibits neural stemness and can convert fibroblasts to neurons through transdifferentiation^50,51^. Furthermore, cluster 3 members tend to highly specifically regulate ArN genes (**Fig. 4E**, **Extended Data Fig. 8A**, ‘Fraction DEGs that are ArN genes’), with POU3F2 alone covering ∼10% of all ArN genes, suggesting that cluster 3 represents molecular mechanisms through which astrocytes promote neuronal maturity. By contrast, cluster 1 perturbations have a mostly negative *in vivo* mimicry score, with NOTCH1, MYCL, and MYC promoter activations among the most dramatic contributors. NOTCH1 is a well-studied factor that maintains cell stemness in the brain, and plays a role in cancer, along with MYCL and MYC^52^. This suggests that cluster 1 members reduce *in vivo* mimicry score by maintaining neuronal immaturity. Indeed, in contrast to cluster 3, whose members most frequently up-regulate Gene Ontology Biological Process (GOBP) terms related to neuron differentiation, axonogenesis and synaptic transmission, cluster 1 members tend to down-regulate these terms but up-regulate protein translation and DNA metabolism, which are hallmarks of activated neural stem cells^53^ (**Fig. 4F**). Furthermore, cluster 1 members tend to have strong but less specific effects on ArN genes (**Fig. 4E**, **Extended Data Fig. 8A**, Log10(1+n. DEGs) and Fraction DEGs that are ArN genes). For example, POU3F2 activation increased the expression of nervous system development, axonogenesis and synaptic genes, while repressing translation genes, whereas MYC activation did the opposite (**Extended Data Fig. 8B**). Therefore, we have identified three groups of perturbations with discrete functions, where cluster 3, which promotes neuronal maturity, counter-balances cluster 1, which promotes cell stemness.

However, we note that CRISPR epigenome editing tends to maintain target gene regulation more consistently than astrocytes at the time of assay (D14) (**Extended Data Fig. 8C**), in contrast to the highly dynamic regulation of neuronal genes by astrocytes (**Fig. 1E**). This could have resulted in synthetic immaturity-maintaining effects of cluster 1 perturbations that do not reflect the true function of astrocytes. In addition, we observed that some members of cluster 2, such as TFAP2E promoter activation and LHX1 fRE repression, can have strong *in vivo* mimicry scores, even though the cluster on average does not (**Fig. 4E**, **Extended Data Fig. 8A**). Notably, all three clusters are composed of perturbations from multiple Perturb-seq screens, indicating that diverse modes of perturbation can achieve similar transcriptional outcomes.

Therefore, we discovered two discrete groups of molecular events with counter-balancing effects, highlighting parallel mechanisms by which astrocytes regulate neuronal maturity and cell identity (**Fig. 1G**). Finally, we observed that 3857 of the 5330 (72.4%) ArN genes are regulated by at least one of these perturbations (excluding positive controls), which act through a total of 68 local target TF genes (**Fig. 4G**). This indicates that our approach successfully identified regulators that mediate the reprogramming of neuronal gene programs by astrocytes.

In summary, joint analysis of both promoters and non-promoter REs of ArN TF genes revealed the *trans* gene regulatory logic that governs the neuronal transcriptional response to astrocytes, with discrete groups of molecular events exerting opposing effects to achieve an overall more mature neuronal transcriptome when dynamically coordinated by astrocytes.

### Gene programs downstream of NAIs are disrupted in autosomal dominant Alzheimer’s Disease and schizophrenia

Given the neurodevelopmental relevance of the transcriptional reprogramming downstream of NAIs, we asked whether it is enriched for genes altered in disease. To enable this comparison, we curated datasets spanning 14 brain disorders from 21 publications, focusing on those with single-cell gene expression-based case/control studies and genetic associations (**Table S17**, **Methods**)^54–76^.

First, we tested whether ArN genes (FDR < 0.01, **Fig. 1D**) and nGMs are enriched for disease genes, considering only protein-coding genes, with all protein-coding genes detectable in our bulk RNA-seq as background (Fisher’s exact test, FDR adjustment). We found several significant associations after multiple comparison corrections, including with familial (but not sporadic) Alzheimer’s disease (AD), autism spectrum disorder (ASD), and schizophrenia (SCZ) (**Fig. 5A**, **Table S18**). There are multiple lines of evidence for the ASD and SCZ associations. For ASD, we found enrichment in genes associated with rare coding variants implicated using whole exome sequencing (‘Fu2022, WES genes’) and genes differentially expressed in bulk brain tissue or single-cell excitatory neurons in case/control studies (‘Gandal2018, bulk RNA-seq, DEGs’ and ‘Velmeshev2019, scRNA-seq, DEGs’). For SCZ, we found enrichment in genes differentially expressed in bulk brain tissue (‘Gandal2018’) and single-cell excitatory neurons (‘Ruzicka2024’) in case/control studies, and the excitatory neuron component of a neuron-astrocyte coordinated gene program that reduced expression in both SCZ and ageing (‘Ling2024’)^68^. Furthermore, these enrichments are primarily found in nGM1, 3, and 4, which are enriched for synaptic pathways, which are thought to be disrupted in ASD and SCZ^59,69^.

**Figure 5:**
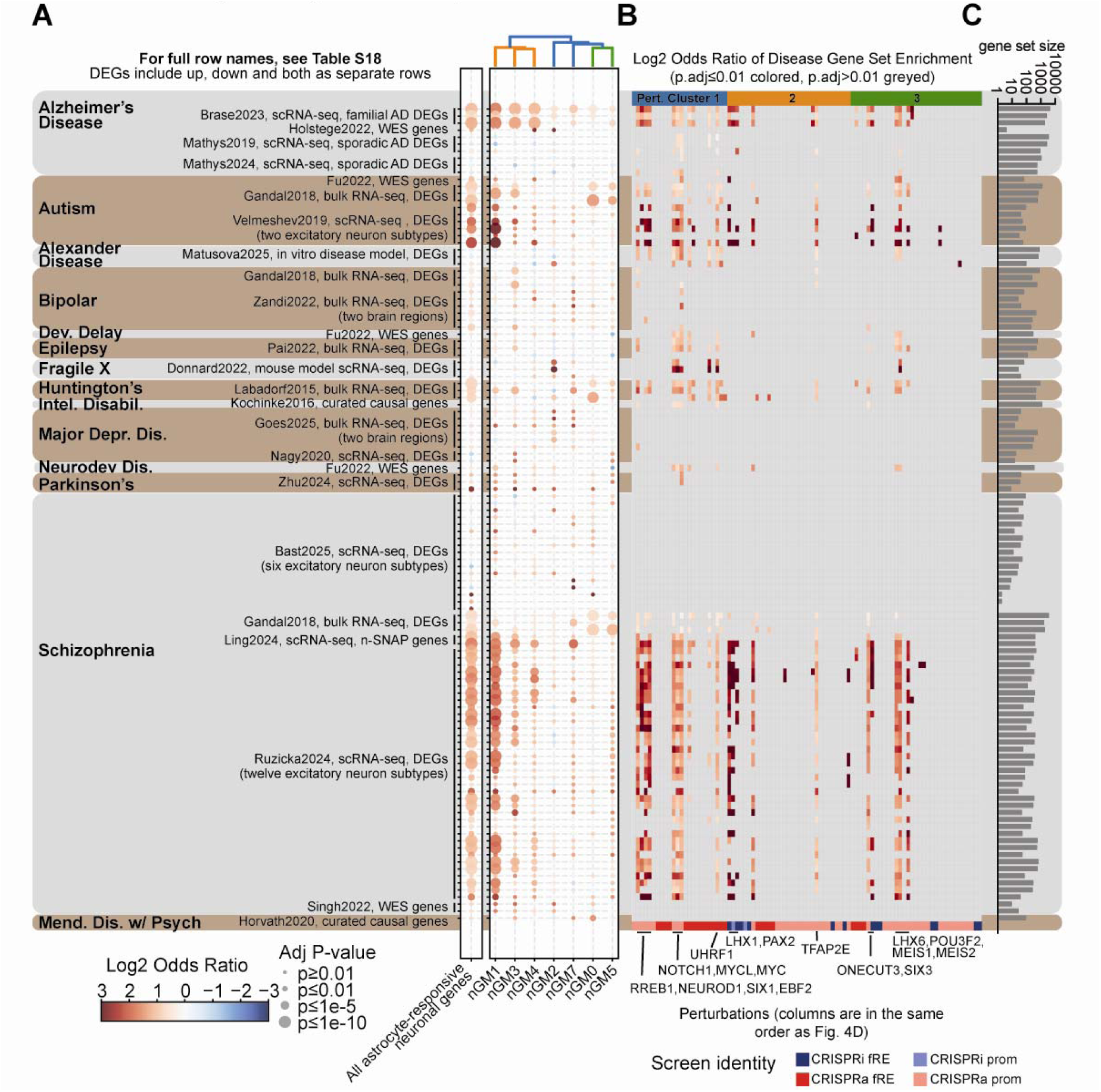
Astrocyte-responsive neuronal gene programs are disrupted in autosomal dominant Alzheimer’s Disease, autism, and schizophrenia. (A) Dotplot showing the enrichment of ArN genes (union set, and individual nGMs, as described in Fig. 1E) for various neurodevelopmental, neuropsychiatric, and neurodegenerative disease or disorder gene sets. Gene sets are labeled with publication (first author last name followed by year of publication), and name or category of the disease/disorder. A detailed list of the gene sets used can be found in **Table S18**. Abbreviations: WES, whole-exome sequencing-based gene set; bipolar, bipolar disorder; dev. delay, developmental delay; intel. disabil., intellectual disability; major depr. dis., major depressive disorder; neurodev. dis., neurodevelopmental disorders; mend. disease w/ psych, Mendelian disease with psychiatric symptoms. (B) Heatmap showing the enrichment of DEGs downstream of TF fRE/promoter perturbations for the same diseases and disorders as shown in (**A**) of this figure. Only statistically significant associations (FDR≤0.01) are colored. (C) Bar chart showing the log10-transformed sizes of the disease gene sets.

In addition, we found associations with disease gene sets for many of the 88 perturbations with strong transcriptional effects described in **Fig. 4D** (**Fig. 5B**). These enrichments are concentrated on ADAD, ASD, and SCZ, mirroring the enrichments found for ArN genes. Perturbations with associations are found across all 3 clusters, suggesting that perturbing genes that make neurons either more or less mature could both contribute to neurological conditions. Genes previously found to drive strong *in vivo* mimicry scores, including *POU3F2*, *LHX1*, and *NOTCH1*, regulate many disease gene sets. This suggests that, in the future, the epigenetic control of these genes could be harnessed for therapeutic purposes.

Next, we tested the same disease datasets against NrA genes (FDR < 0.01, **Extended Data Fig. 2B**) and aGMs. We found both overlapping and unique associations relative to ArN genes (**Extended Data Fig. 9A**, **Table S18**). Like ArN genes, NrA genes are also associated with ADAD and SCZ, suggesting that these conditions converge on the disruption of NAIs that manifests in both neurons and astrocytes. However, ArN genes and NrA genes are largely non-overlapping (**Extended Data Fig. 9C**), suggesting that gene dysregulation in these conditions manifests distinctly in neurons and astrocytes. The association of NrA genes with SCZ is also supported by both cases/controls-based gene expression analysis (‘Gandal2018’ and ‘Ruzicka2024’) and the neuron-astrocyte gene program identified by latent factor analysis (‘Ling2024’)^68^. Interestingly, Ling and colleagues showed that the SCZ-associated latent factor (which they termed synaptic neuron and astrocyte program, or SNAP) is driven by both neurons and astrocytes, but through distinct sets of genes (**Extended Data Fig. 9D**). Consistently, we observed that ArN genes and NrA genes are respectively enriched for neuron- and astrocyte-driver genes of SNAP, indicating a convergence between our findings and theirs. This suggests that the disruption of NAIs leads to the dysregulation of the SCZ-associated gene programs in both neurons and astrocytes. Furthermore, NrA genes are enriched for genes altered in sporadic AD, epilepsy, Fragile X syndrome, Huntington disease and Parkinson’s disease (**Extended Data Fig. 9A**). The association of genes dysregulated in both sporadic and familial AD genes with NrA genes supports that they converge on gene dysregulation in astrocytes. Moreover, NrA genes strikingly include 5 of 6 AD causal genes identified through rare variants (‘Holstage2022 WES genes’), indicating that AD-causal genes are responsive to NAIs. These results together suggest that both common and rare AD variants converge on NAI disruption and its subsequent effects on astrocytes.

In summary, we found that gene programs downstream of NAIs are altered in neurological diseases and disorders, with genes in both neurons and astrocytes dysregulated in familial AD and SCZ, suggesting that the disruption of NAIs contributes to these conditions. Gene regulatory networks downstream of epigenetic perturbations overlap with disease genes, and thus might be further exploited for disease treatment.

### ArN TF perturbations reprogram neuronal electrophysiology and neurite morphology

Given the strong transcriptional influence of ArN TFs on genes involved in neuronal differentiation, axonogenesis, and synaptogenesis, we investigated their impact on neuronal electrophysiology and neurite morphology. We emphasized CRISPRa promoter perturbations, which span all three perturbation clusters (**Fig. 4D**), and selected 11 for functional characterization. We further added three CRISPRi fRE perturbations (**Table S19**). Moreover, we included two non-targeting sgRNAs (sgNT1 and sgNT2) as well as neurons co-cultured with astrocytes as controls.

We used time-course multi-electrode arrays (MEA) recording to measure neuronal electrophysiology in cultured neurons for up to five weeks (**Fig. 6A**). Neurons with non-targeting control gRNAs began to spontaneously fire after a week of maturation that increased throughout the course of the experiment (**Extended Data Fig. 10A**, **Fig. 6B**, sgNT2). As expected, astrocytes by themselves did not fire (**Extended Data Fig. 10B**). Co-culture with astrocytes increased firing rates in both the CRISPRa and CRISPRi cell lines starting at week three or four, consistent with previous observations (**Fig. 6C**, **Extended Data Fig. 10C**) ^21^. Three of the CRISPRa perturbations (*OLIG3*, *POU3F2*, and *MEIS2*) increased neuronal firing rate within the same time window in the absence of astrocytes, indicating that they at least partially mimic astrocyte presence to reprogram neuronal electrophysiology.

**Figure 6:**
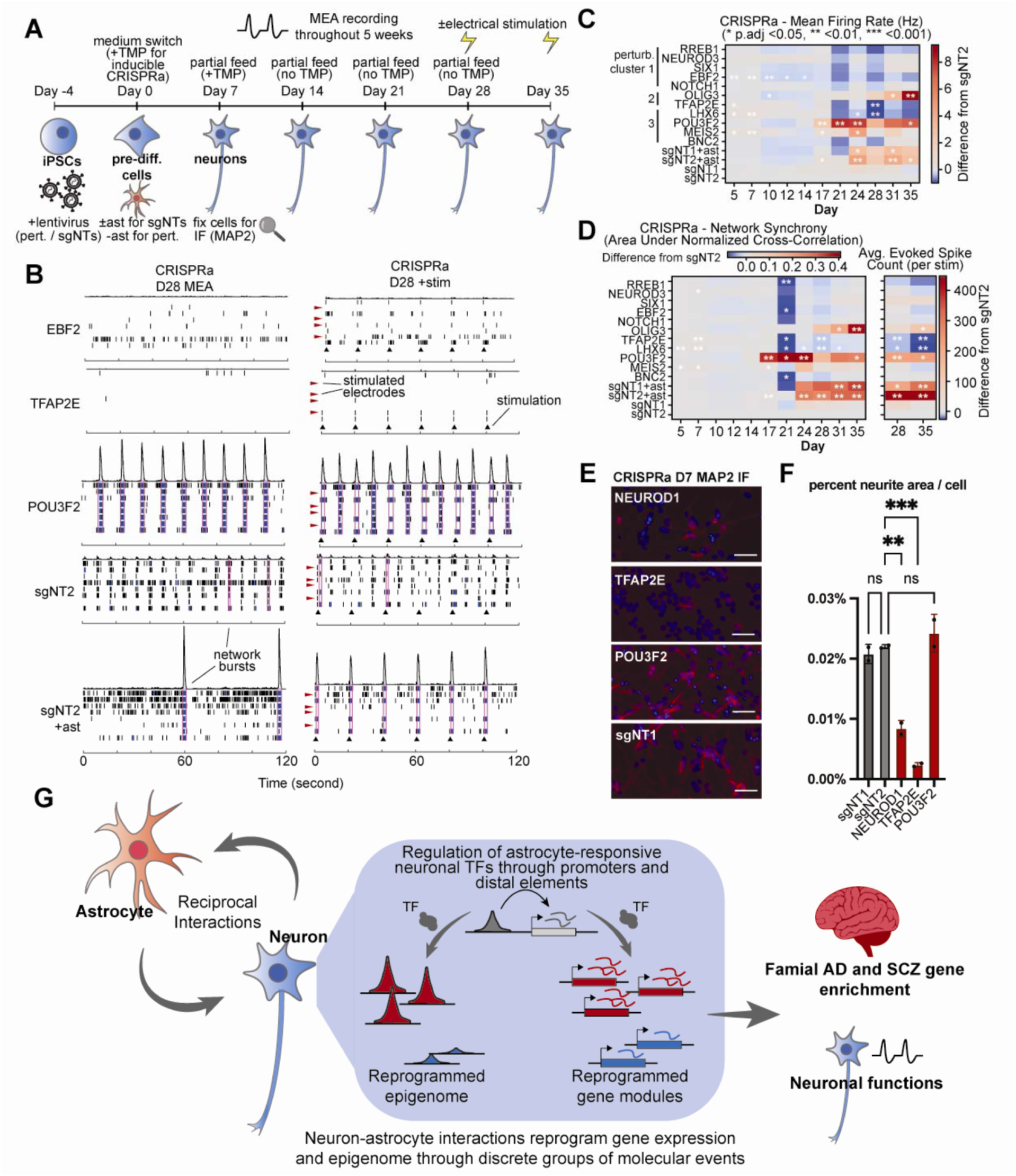
ArN TF perturbations reprogram neuronal electrophysiology and neurite morphology. (A) Schematic showing the MEA and IF experimental setup. (B) Raster plots showing the neurons’ firings on day 28, with or without electrical stimulation, across 8 electrodes of a representative well of 3 example CRISPRa perturbations, one negative control (sgNT2) and negative control co-cultured with astrocytes. Purple boxes indicate network bursts. Black arrows indicate electrical stimulations. Red arrows indicate the 4 electrodes (out of 8) that were stimulated, whose activity was filtered out within a short time window after stimulation to remove artifacts. (C) Heatmap showing relative increase or decrease over negative control (sgNT2) of firing rate (Hz) of neurons with 11 individual CRISPRa perturbations, negative control neurons, or neurons co-cultured with astrocytes. 12 replicates per condition were analyzed. (D) Heatmap showing relative increase or decrease over negative control (sgNT2) of network synchrony (area under normalized cross-correlation) and average evoked spike count per stimulation of neurons with 11 individual CRISPRa perturbations, negative control neurons, or neurons co-cultured with astrocytes. 12 replicates per condition were analyzed. (E) Immunofluorescence images of the neuronal dendritic marker MAP2 across 3 example CRISPRa perturbations and negative control (sgNT1). (F) Quantification of neurite area per cell as a percentage of the entire image. Two images per condition were analyzed. **, adjusted p < 0.01; ***, adjusted p < 0.001. (G) A cartoon summary of the findings from this study.

In addition, co-culturing with astrocytes induced robust network bursts by day 28 that can be paced to low frequency (0.05 Hz) electrical stimulation, indicative of synchronous and functionally mature neuronal networks, as previously observed^22,77^ (**Fig. 6B, D**, **Extended Data Fig. 10D**). CRISPRa perturbations of OLIG3 and POU3F2 enhanced network synchrony and response to stimulation, supporting their ability to increase neuronal maturity, consistent with the interpretation above of POU3F2 activation’s transcriptional effects (**Fig. 4E**). In contrast, TFAP2E, a perturbation with strongly negative *in vivo* mimicry score, despite increasing firing rates early on, decreased firing rate, network synchrony, and response to stimulation in the long term (**Extended Data Fig. 10A**, **Fig. 6B, D**). Notably, none of the five perturbations from cluster 1 increased these measurements, consistent with our interpretation that cluster 1 represents perturbations that decrease neuronal maturity (for example, EBF2 in **Extended Data Fig. 10A**, **Fig. 6BD**). By MAP2 immunofluorescence staining, we observed that NEUROD1 (a cluster 1 perturbation) and TFAP2E strikingly reduced dendrite outgrowth, while POU3F2 maintained it (**Fig. 6E, F**), consistent with both transcriptional and electrophysiological data.

In summary, both astrocyte co-culture and the synthetic epigenetic perturbations that mimic co-culture can reprogram neuronal electrophysiology and neurite morphology.

## Discussion

Though widely appreciated to be critical components of organismal development, tissue homeostasis, and disease progression, the epigenetic consequences of heterotypic cell-cell interactions have not been finely or comprehensively mapped. Using a neuron-astrocyte co-culture model, we gained insights into how the genome’s function is reprogrammed by heterotypic cell-cell interactions (**Fig. 6G**) by measuring their impact on gene expression and chromatin landscape, as well as by Perturb-seq experiments that recapitulate the individual transcriptional and epigenetic events downstream of NAIs in mono-cultured neurons.

To investigate the genomic effects of NAIs, we performed bulk RNA-seq and ATAC-seq and showed that the neuronal genome is extensively reprogrammed by NAIs, affecting about a quarter of expressed genes and ∼10% of the open chromatin regions. The scale of thousands of responsive genes is consistent with previous reports^19,20^, but pairwise overlap among the three studies is moderate (∼1.5x enrichment over chance for all three pairs), suggesting that choice of the model system influences the observed neuronal response to glia. Specifically, the key differences include that 1) Burke et al. used a medium-induced differentiation method rather than Ngn2 activation; 2) Pietilainen et al. used a mixed population of glia, whereas our study and Burke et al. used purified mouse and rat astrocytes, respectively. Nevertheless, 303 genes were found to be common to all three studies, with enriched GO terms dominated by synapse-and nervous system development-related genes (**Extended Data Fig. 1E**), supporting the notion that glial cells play central roles in the regulation of neuronal differentiation and synaptic function. Furthermore, we showed that in our co-culture model, neurons have transcriptomes that are more strongly correlated to *in vivo* neurons compared to mono-cultured neurons, as well as form more functional networks, indicating they are more mature and better models of physiological processes. Neurological and neuropsychiatric disorders such as epilepsy, Rett Syndrome, and schizophrenia manifest in altered network activity in iPSC-derived neuronal models, suggesting that a more functional network would improve disease modeling and drug screening for these disorders^78–80^. Consistently, astrocyte co-culture has been reported to improve drug screening of neurotoxicity^81^. Importantly, we discovered factors downstream of NAIs, such as *POU3F2*, that enhance network activity, which could be harnessed to improve network functions of current iPSC-derived neuronal models for these applications.

Astrocytes have been increasingly recognized as active drivers of and therapeutic targets in neurological pathologies, particularly in neurodegenerative diseases such as AD^82,83^. Thus, studying the molecular consequences of NAIs could lead to insights in disease mechanisms and new therapeutic targets. We found that both ArN and NrA gene programs are commonly altered in familial AD and SCZ, confirming the relevance of NAIs in these conditions. Consistently, iPSC-derived astrocytes harboring familial AD mutations exhibit a wide range of AD hallmarks, such as increased β-amyloid production and altered cytokine release^84,85^. In addition, important pathways at the neuron-astrocyte interface, including glutamate uptake and lactate secretion, are impaired, supporting the disruption of NAIs in familial AD. On the other hand, although GWAS studies have mostly implicated neurons, not astrocytes, in the genetics of SCZ risk^75,86^, astrocyte genes could nevertheless be dysregulated through the disruption of NAIs. In addition, transplantation of SCZ patient iPSC-derived glial progenitor cells and astrocytes into mice suggested abnormality in astrocyte differentiation and function in SCZ^87,88^. Therefore, neuron-astrocyte co-culture can serve as a powerful model to parse out the cell-autonomous and non-cell-autonomous genetic drivers of epigenetic and transcriptional dysregulation in familial AD and SCZ.

To illuminate the *cis* and *trans* gene regulatory mechanisms that govern the response of the neuronal genome to astrocytes, we designed CRISPRi and CRISPRa Perturb-seq screens targeting the promoters and non-promoter REs of ∼200 ArN TF genes, with the goal of recapitulating the impact of NAIs one molecular event at a time. First, we discovered functional REs for ∼50 of the TF genes, revealing the often complex *cis* regulatory landscapes and expanding the catalog of experimentally established REs in human neurons that can be harnessed in the future to engineer gene expression for applications such as *cis*-regulation therapy^89^. Second, we discover that, strikingly, epigenome editing events that mimic the influence of astrocytes form discrete groups that exert opposing effects on the transcriptome. Importantly, these TFs also reprogram relevant neuronal functions, albeit not perfectly. For example, *LHX6* activation decreased electrophysiological function while being a member of cluster 3 that was predicted to increase neuronal maturity. Our study presents a step toward understanding how genomic information propagates between neurons and astrocytes, providing a blueprint for applying genomics and CRISPR perturbation approaches to uncover the links between cell environment and TF genes, gene modules, and cellular functions.

We note that even with our comprehensive perturbation strategies, we did not identify regulators of all ArN genes identified in bulk RNA-seq (**Fig. 4G**). This hints at future opportunities to improve perturbation strategies to better mimic astrocyte-mediated effects on neurons. Firstly, we selected TF gene targets based on mRNA-level response to astrocytes (**Extended Data Fig. 5A**), but TF proteins are often regulated at the protein level^90^. Therefore, selecting additional targets based on, e.g., motif enrichment in the ArN ATAC-seq peaks might lead to a more complete view of the gene regulatory networks. Secondly, since TFs often interact, future combinatorial TF perturbations could lead to better mimicry of astrocyte-mediated effects in a non-additive manner. Thirdly, we showed that astrocyte-mediated gene expression reprogramming is highly temporally dynamic (**Fig. 1E**), while the CRISPRi/a approach we used was constitutive. Future studies using temporally controlled perturbation strategies might allow more physiological reprogramming of the gene modules.

We acknowledge the limitations of the co-culture model of *Ngn2*-induced hiPSC-derived neurons and primary mouse astrocytes. First, *Ngn2* induction bypasses natural intermediate progenitor states, and thus does not fully model in vivo neurodevelopment ^23^. Furthermore, there are well-documented differences between human and mouse astrocytes, including morphology, gene expression, and function ^91^. Therefore, the results described here should be interpreted as a significant step towards, but not the comprehensive description of how neuronal genome function responds to astrocytes.

Altogether, these discoveries highlight the importance of NAIs in the regulation of neurodevelopment- and disease-relevant gene modules, serving as a foundation toward the understanding of how genome function and therefore genetic variation can be interpreted in lieu of the microenvironment. We expect this work will inspire future studies investigating how disease-causal genetic variants, such as for familial AD, in one cell type might impact neighboring cell types, and the underlying intercellular and intracellular molecular mechanisms that mediate the reprogramming of genome function by cell-cell interactions in diverse biological systems.

## Data Availability

- [IGVF data portal accession numbers]

- COMING SOON. DATA SUBMISSION IN PROGRESS.

## Code Availability

The custom code and scripts used for all analyses and figure generation are available in the GitHub repository: https://github.com/liboxun/Li-et-al-2026-Neuron-astrocyte-interactions. Upload is expected to complete by the time of formal publication.

## Acknowledgements

The authors thank Dr. Yihan Wang for help and advice with pySpade; Dr. Yin Shen and Dr. Martin Kampmann (UCSF) for WTC11 iPSC cell line with dCas9-KRAB and dCas9-VPH, respectively; Dr. Scott Soderling (Duke University) for access to Axion Maestro Pro; Duke Sequencing and Genomic Technologies core facility access to next-generation sequencing equipment; the members of the Gersbach, Crawford, and Eroglu laboratories for comments and suggestions throughout this work. This work was supported by NIH grants UM1-HG012053, R01-MH125236, UM1-HG012003, UM1-HG012051, and RM1-HG011123; NSF EFMA-1830957 (C.A.G.), and Open Philanthropy (C.A.G.).

## Author contributions

**Boxun Li**: Conceptualization, Formal analysis, Investigation, Methodology, Software, Validation, Visualization, Writing – Original Draft, Writing – Review & Editing. **Kevin T. Hagy**: Formal analysis, Investigation, Validation, Writing – Original Draft. **Alexias Safi**: Investigation, Writing – Original Draft. **Michael A. Beer**: Formal analysis, Methodology, Software, Writing – Original Draft, Writing – Review & Editing. **Alejandro Barrera**: Formal analysis, Visualization, Software, Writing – Original Draft. **Sara Geraghty**: Formal analysis, Software, Writing – Review & Editing. **Ruhi Rai**: Visualization. **Alyssa N. Pederson**: Investigation. **Samuel J. Reisman**: Investigation, Writing – Review & Editing. **Michael I. Love**: Methodology, Writing – Review & Editing. **Patrick F. Sullivan**: Methodology, Writing – Review & Editing. **Cagla Eroglu**: Conceptualization, Methodology, Writing – Review & Editing. **Gregory E. Crawford**: Funding acquisition, Supervision, Writing – Review & Editing. **Charles A. Gersbach**: Conceptualization, Funding acquisition, Methodology, Supervision, Writing – Review & Editing

## Competing interests

C.A.G. is a co-founder of Tune Therapeutics, Sollus Therapeutics, and Locus Biosciences and is an advisor to Tune Therapeutics, Sollus Therapeutics, Pappas Capital, and Sarepta Therapeutics. G.E.C., and C.A.G. are inventors on patents or patent applications related to CRISPR epigenome editing and screening technologies. P.F.S. was a consultant and shareholder for Neumora Therapeutics.

## Extended Data Figures

**Extended Data Figure 1:**
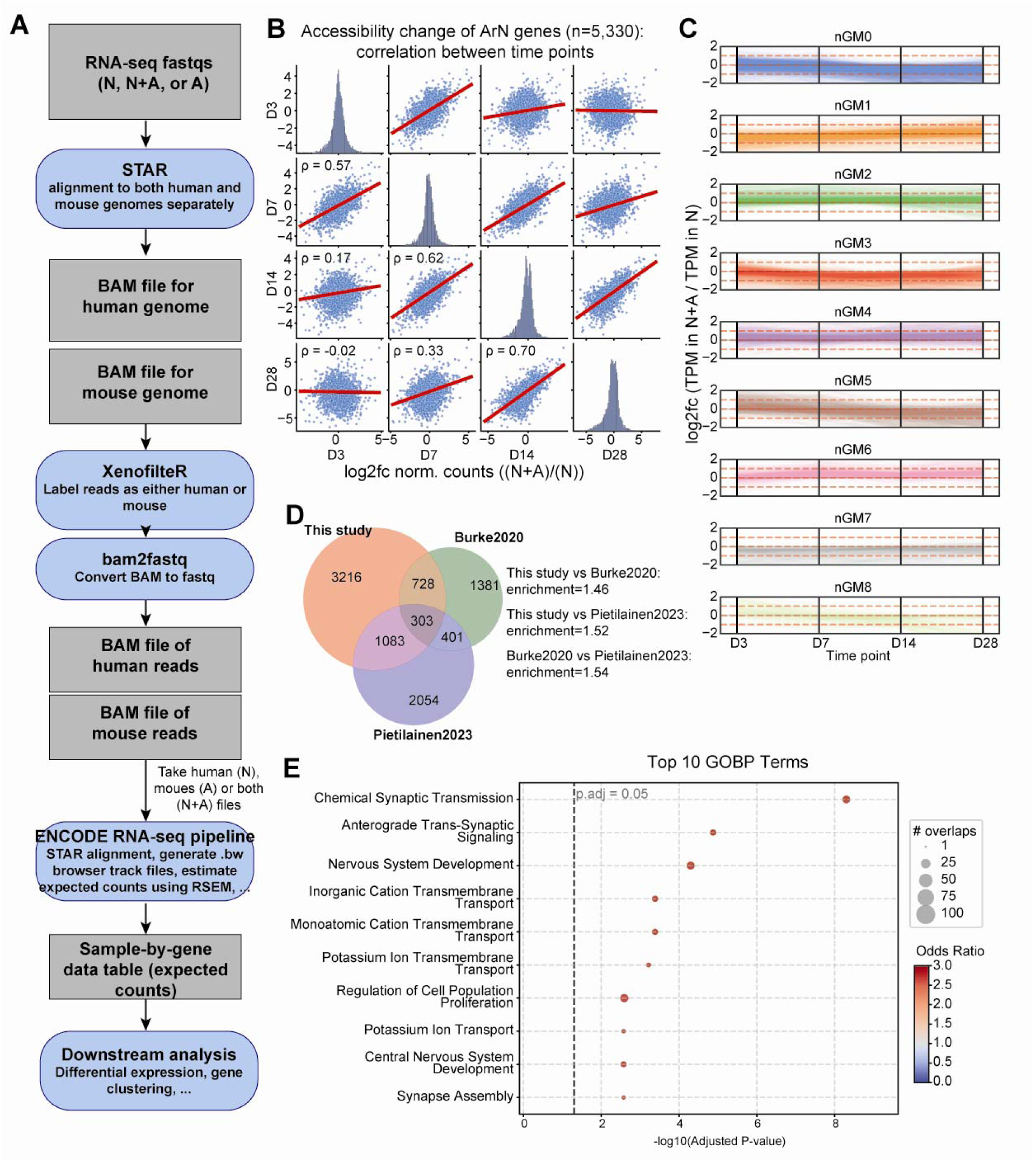
Computational pipeline and additional results related to Figure 1. (A) Diagram showing the bulk RNA-seq analysis pipeline. (B) Scatterplots showing the the log2fc(normalized counts) changes between (N+A) and N of ArN genes between pairs of time points. (C) Parallel plots showing the different trends of the log2fc(normalized counts) changes of ArN gene programs (nGMs) between N+A and N. (D) Venn diagram showing the overlaps between the genes found to be implicated downstream of neuron-glia interactions from this study, Burke et al., 2020, and Pietilainen et al., 2023. (E) Dotplot of top 10 enriched GOBP terms of the genes in 3-way overlap among this study, Burke et al., 2020, and Pietilainen et al., 2023.

**Extended Data Figure 2:**
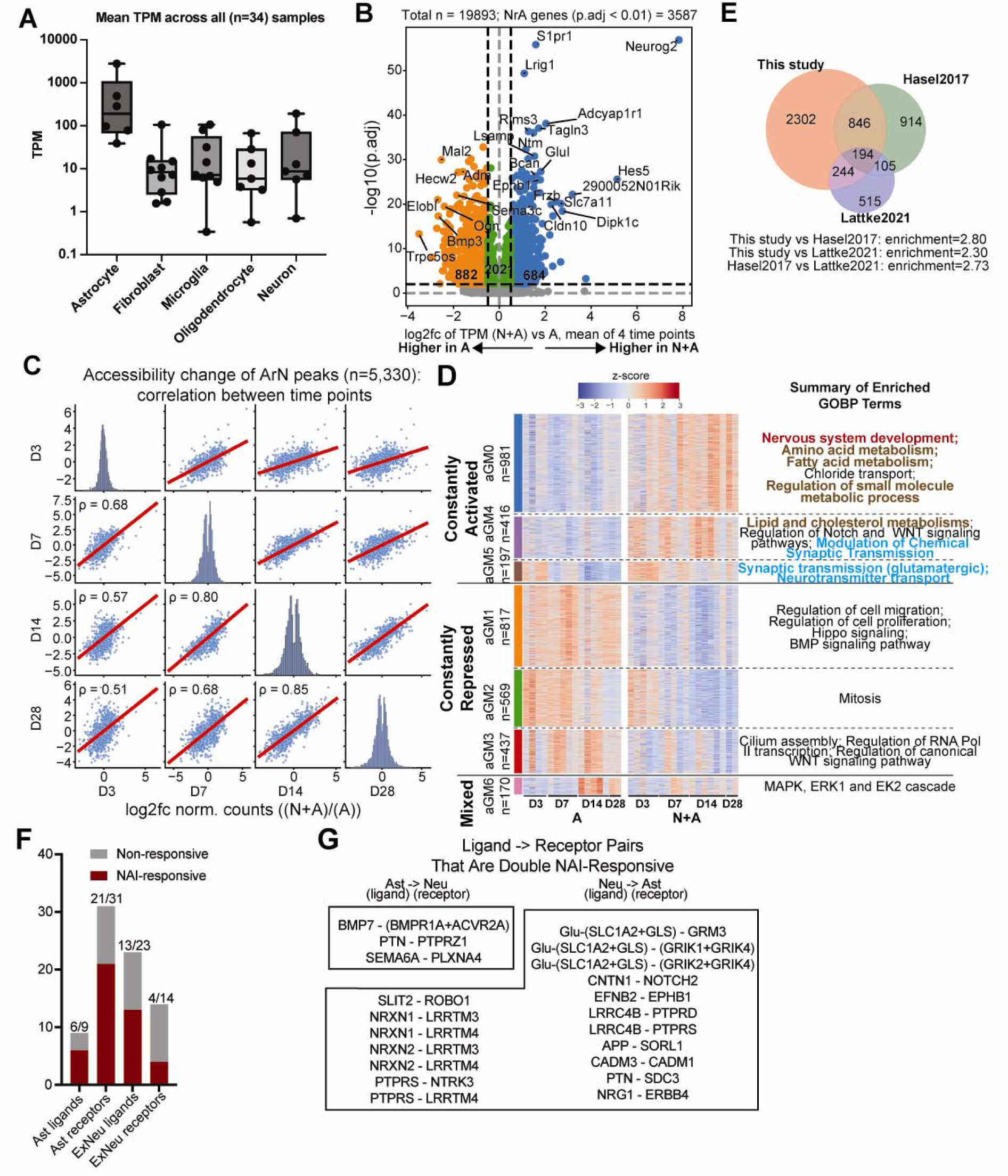
Neuron-astrocyte interactions reprogram astrocytic gene expression programs. (A) Boxplot showing the expression of marker genes of different cell types in our astrocyte transcriptomes (both from mono- and co-culture conditions) to confirm high astrocytic purity. (B) Volcano plot showing the changes of the NrA genes between N and N+A conditions, average of all 4 time points. (C) Scatterplots showing the log2fc(normalized counts) changes of between (N+A) and N of the NrA genes between pairs of time points. (D) Heatmap showing z-scaled expression of the NrA genes across mono- and co-culture conditions, time points, and replicates, with enriched GOBP terms manually summarized. (E) Venn diagram showing the overlaps between the genes found to be implicated downstream of neuron-glia interactions from this study, Hasel et al., 2017, and Lattke et al., 2021. (F) Barplot showing the numbers of receptor or ligand genes inferred by CellChat to mediate neuron-astrocyte interactions that are responsive to neuron-astrocyte interactions in our RNA-seq analysis (NAIs). (G) Lists of receptor-ligand pairs that are double-responsive to NAIs in our RNA-seq analysis.

**Extended Data Figure 3:**
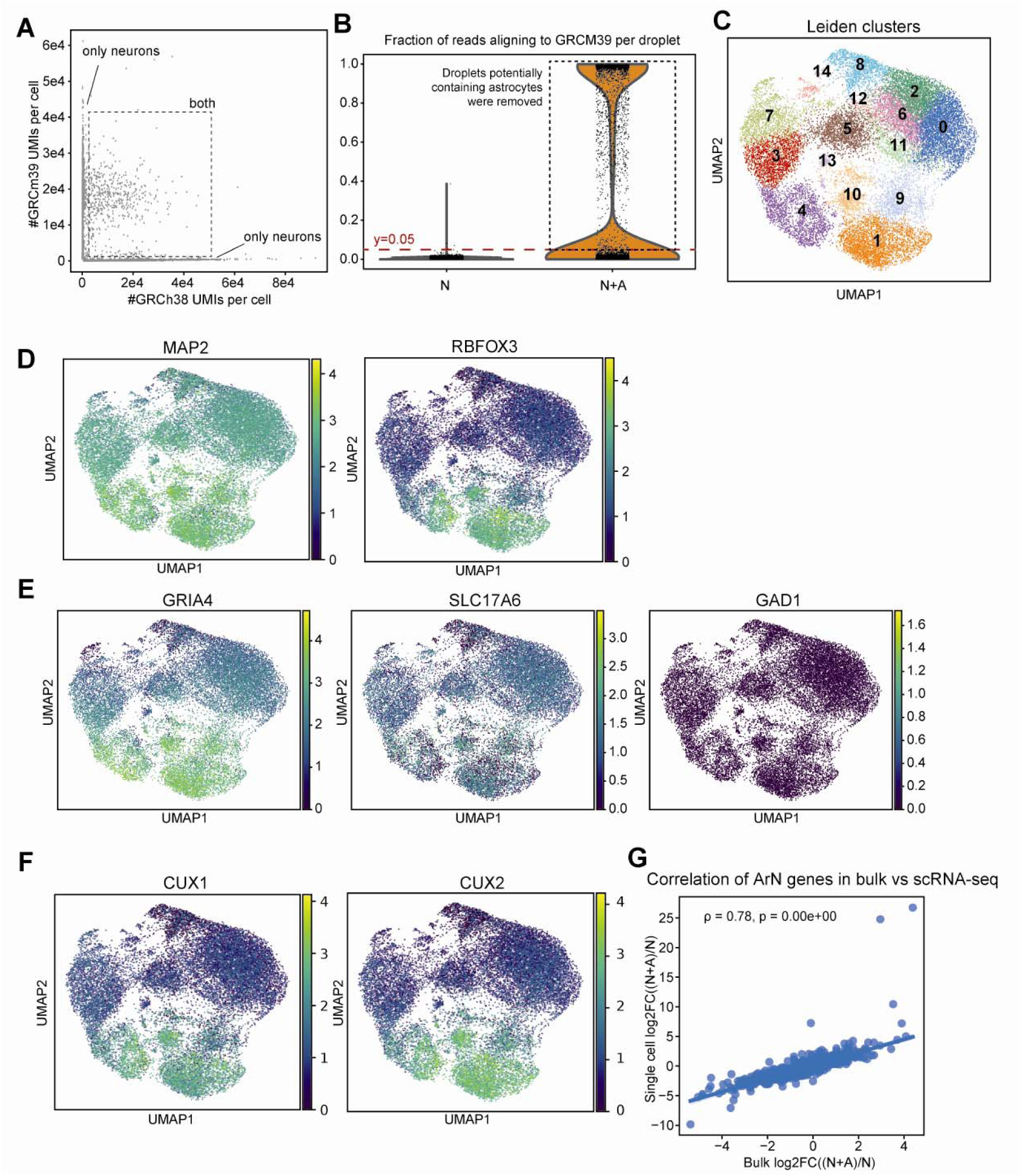
Single-cell RNA-seq validation of ArN genes. (A) Scatter plot showing numbers of reads aligning to the human (GRCh38) and mouse (GRCm39) genomes, respectively, per droplet from the N+A 10X Genomics scRNA-seq lane. (B) Violin showing the fraction of reads aligning to the mouse (GRCm39) genome per droplet from the N and N+A 10X Genomics scRNA-seq lanes, respectively. Fraction < 0.05 was used to filter out any droplets potentially containing astrocytes in the N+A lane. (C) UMAP plot of the N vs N+A scRNA-seq data showing cells colored by Leiden cluster identity. (D) UMAP plots showing single-cell expression of pan-neuronal markers MAP2 and RBFOX3. (E) UMAP plots showing single-cell expression of glutamatergic (GRIA4, SLC17A6) and GABAergic (GAD1) markers. (F) UMAP plots showing single-cell expression of cortical neuronal marker CUX1 and CUX2. (G) Scatterplot showing the Pearson correlation between bulk and single-cell log2FC of the 6,549 ArN genes identified in the N vs N+A scRNA-seq dataset.

**Extended Data Figure 4:**
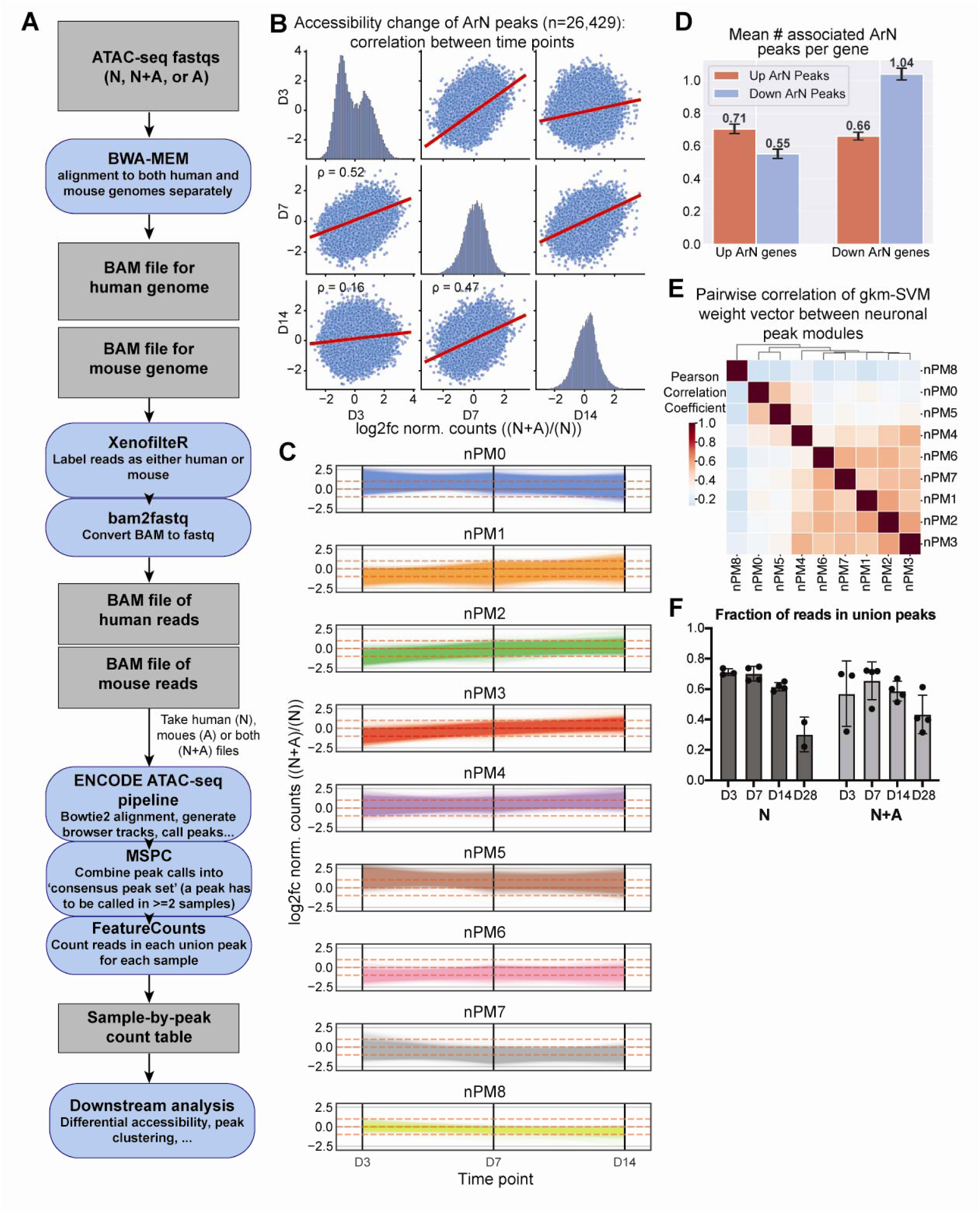
Computational pipeline and additional results related to Figure 2. (A) Diagram showing the bulk ATAC-seq analysis pipeline. (B) Scatterplots showing the log2fc(CPM) changes between (N+A) and N of ArN peaks between pairs of time points. (C) Parallel plots showing the different trends of the log2fc(CPM) changes of ArN peak programs (nPMs) between N+A and N. (D) Barplots showing the average numbers of up and down ArN peaks per ArN gene. Error bars: SEM. (E) Pairwise correlation between nPMs based on gkm-SVM (‘motif’) weight vectors. (F) Barplot showing the fraction of reads in union peaks, a key quality metric, across time points and biological replicates.

**Extended Data Figure 5:**
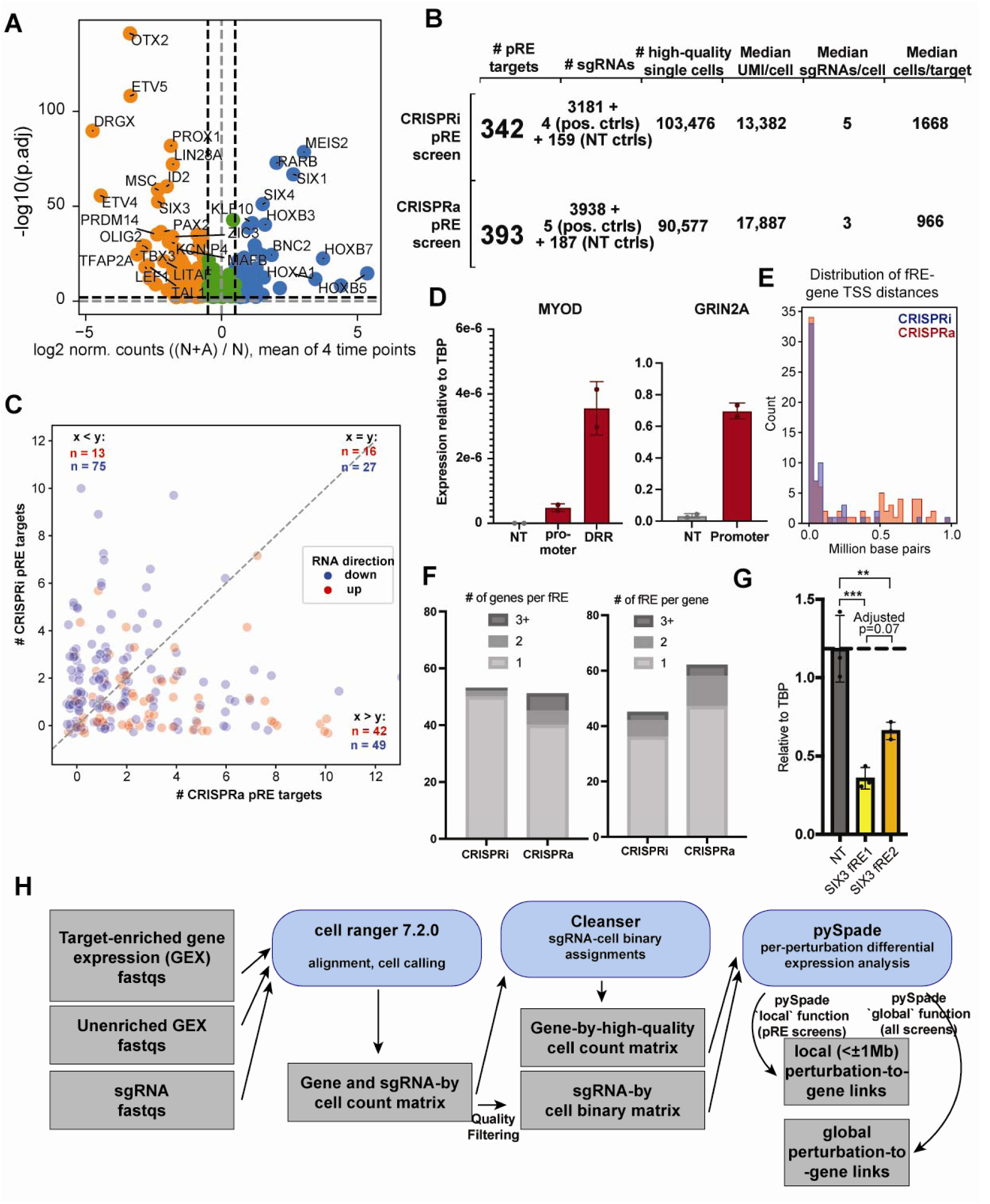
pRE screens local analysis. (A) Volcano plot showing the changes of n=222 ArN TF genes selected for Perturb-seq between N and N+A conditions, average of 4 time points. (B) Table showing statistics of sgRNA library design and data quality metrics for CRISPRi and CRISPRa pRE screens. (C) Scatterplot of showing the numbers of CRISPRi vs CRISPRa pRE targets per ArN TF gene, with genes that are up-regulated by astrocytes at the mRNA level colored in red, and genes down-regulated by astrocytes colored by blue. (D) Barplot showing that the dCas9-p300 cell line activates GRIN2A and MYOD genes by sgRNAs targeted at promoters and a distal regulatory element (^4^) in differentiated neurons. (E) Histogram showing the distributions of base-pair distance between functional RE and target genes identified in the CRISPRi and CRISPRa pRE screens, respectively. (F) Barplots showing the distribution of # of target genes assigned to a functional RE, and # of functional REs assigned to a target gene, respectively. (G) Barplot showing the effects of CRISPRi on two different functional REs of the TF gene SIX3. The statistical test was done by one-way ANOVA followed by Tukey’s multiple comparisons test. **: adjusted p<0.01; ***: adjusted p<0.001. (H) Schematics showing a high-level overview of the single-cell analysis pipeline of the Perturb-seq screens, highlighting the methods used.

**Extended Data Figure 6:**
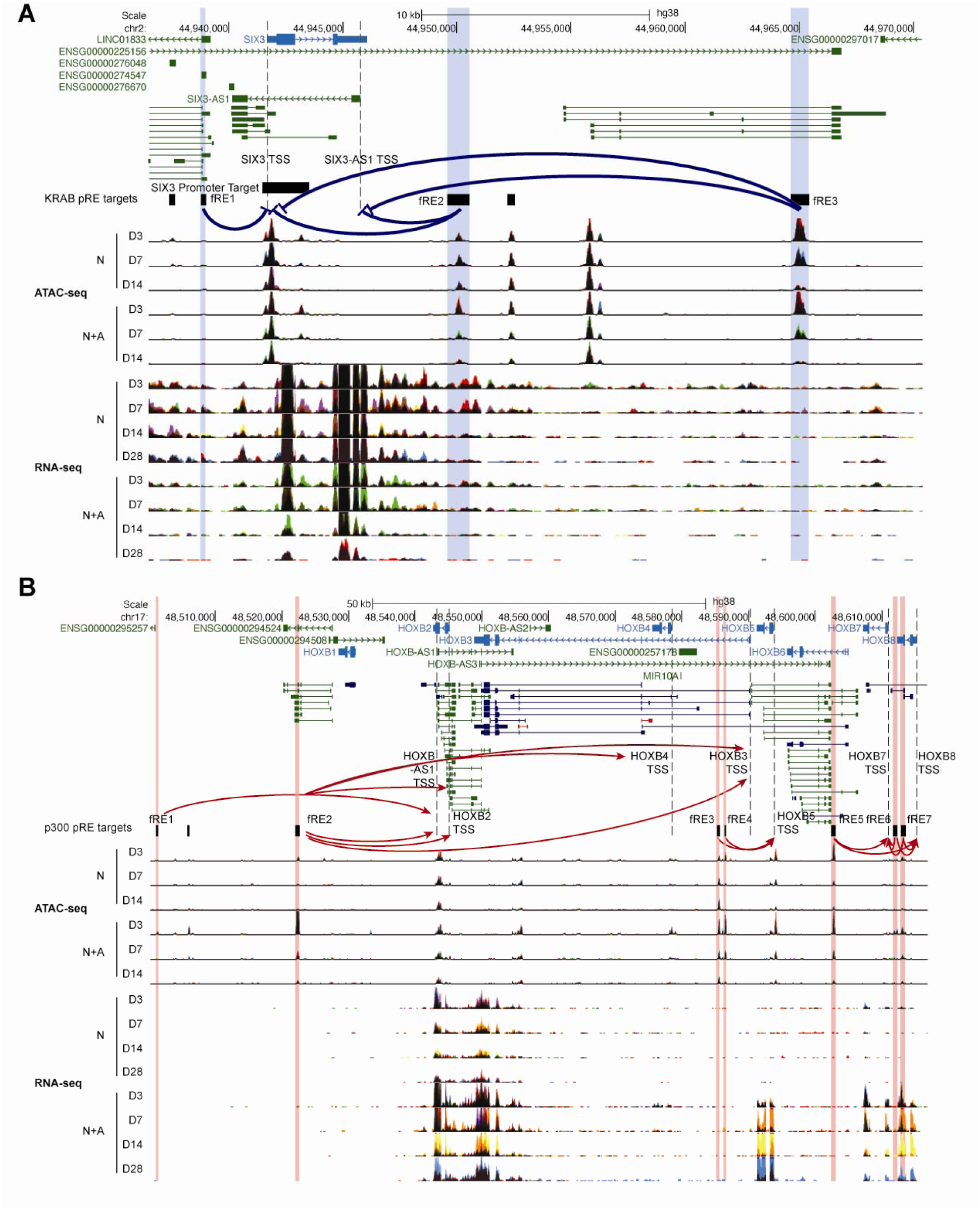
Example loci for the CRISPRi (SIX3) and CRISPRa (HOXB) pRE screens. (A) Genome browser snapshot of the SIX3 gene locus. Blue shade: fREs discovered in the CRISPRi screen. Blue arrows: CRISPRi fRE-gene links. (B) Genome browser snapshot of the HOXB gene locus. Red shade: fREs discovered in the CRISPRa screen. Red arrows: CRISPRa fRE-gene links.

**Extended Data Figure 7:**
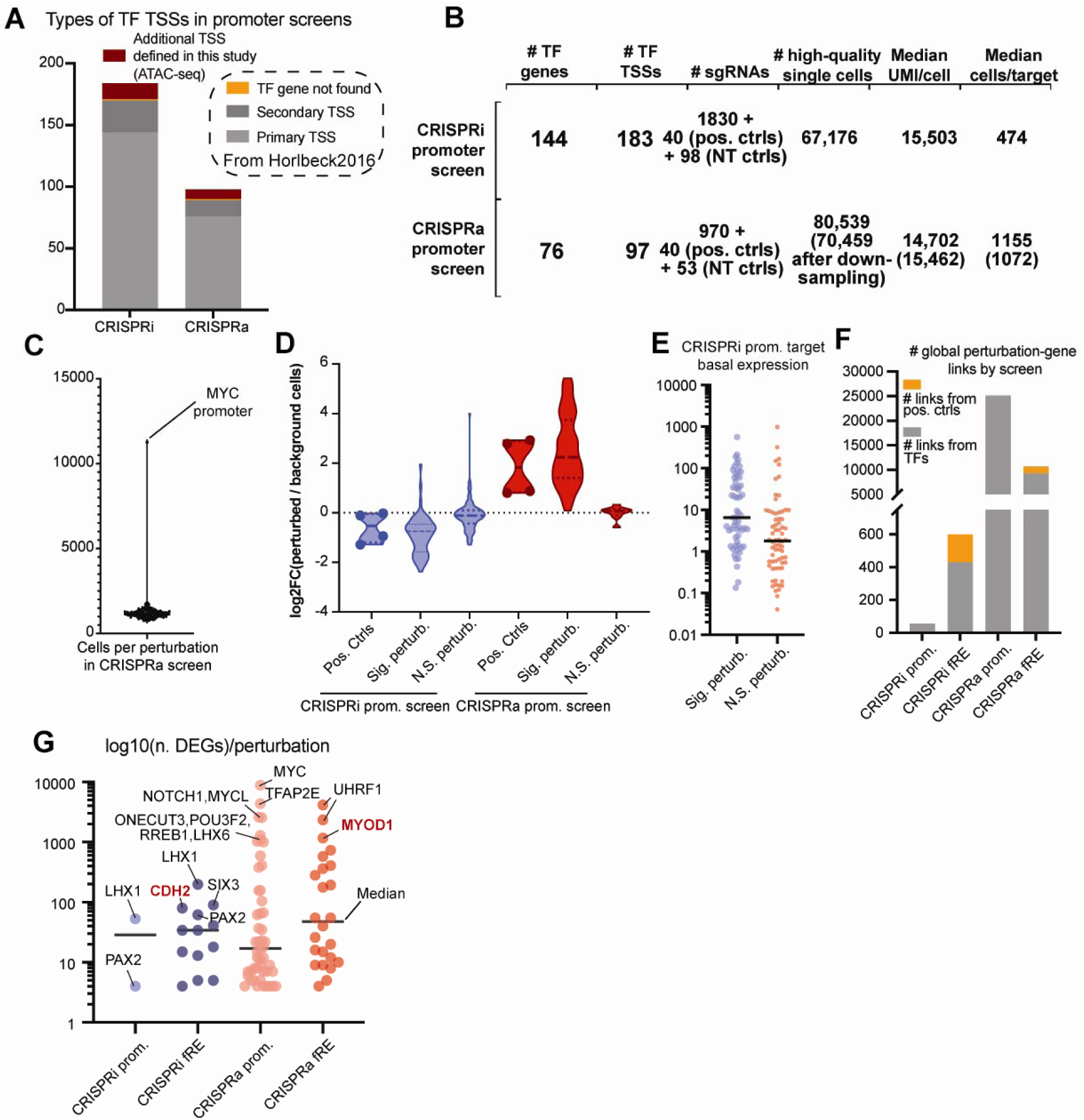
Statistics for the promoter Perturb-seq experiments. (A) Stacked bar plots showing the numbers of targeted annotated primary TSSs, alternative TSSs, and unannotated TSSs based on the RNA-seq data generated in this study, in the CRISPRi/a promoter Perturb-seq experiments, respectively. (B) Table showing statistics of sgRNA library design and data quality metrics for CRISPRi and CRISPRa promoter screens. (C) Violin plot showing the number of cells per target region in the CRISPRa promoter Perturb-seq experiment, highlighting the MYC promoter as an outlier. (D) Violin plot showing the log2FC of the direct target genes in the CRISPRi and CRISPRa promoter Perturb-seq experiments, respectively, separated by positive controls (regardless of significance), TF genes with significant alteration in expression level (Sig. perturb.; p < 0.05), and TF genes with non-significant alteration (N.S. perturb.; p >= 0.05). (E) Jitter plot showing the basal expression levels of CRISPRi promoter target genes that were successfully (‘success’) or unsuccessfully (‘fail’) repressed (F) Bar plot showing the numbers of total global perturbation-gene links discovered by the 4 Perturb-seq experiments. Only those from the positive controls, statistically significant promoters (for the promoter experiments), and fREs (for the pRE experiments) are shown. (G) Jitter plot showing the numbers of global perturbation-gene links per perturbation. Only those from the positive controls, statistically significant promoters (for the promoter experiments), and fREs (for the pRE experiments) are shown. Top perturbations are labeled by its local or direct target gene, for fRE and promoter, respectively. Red bolded are positive controls.

**Extended Data Figure 8:**
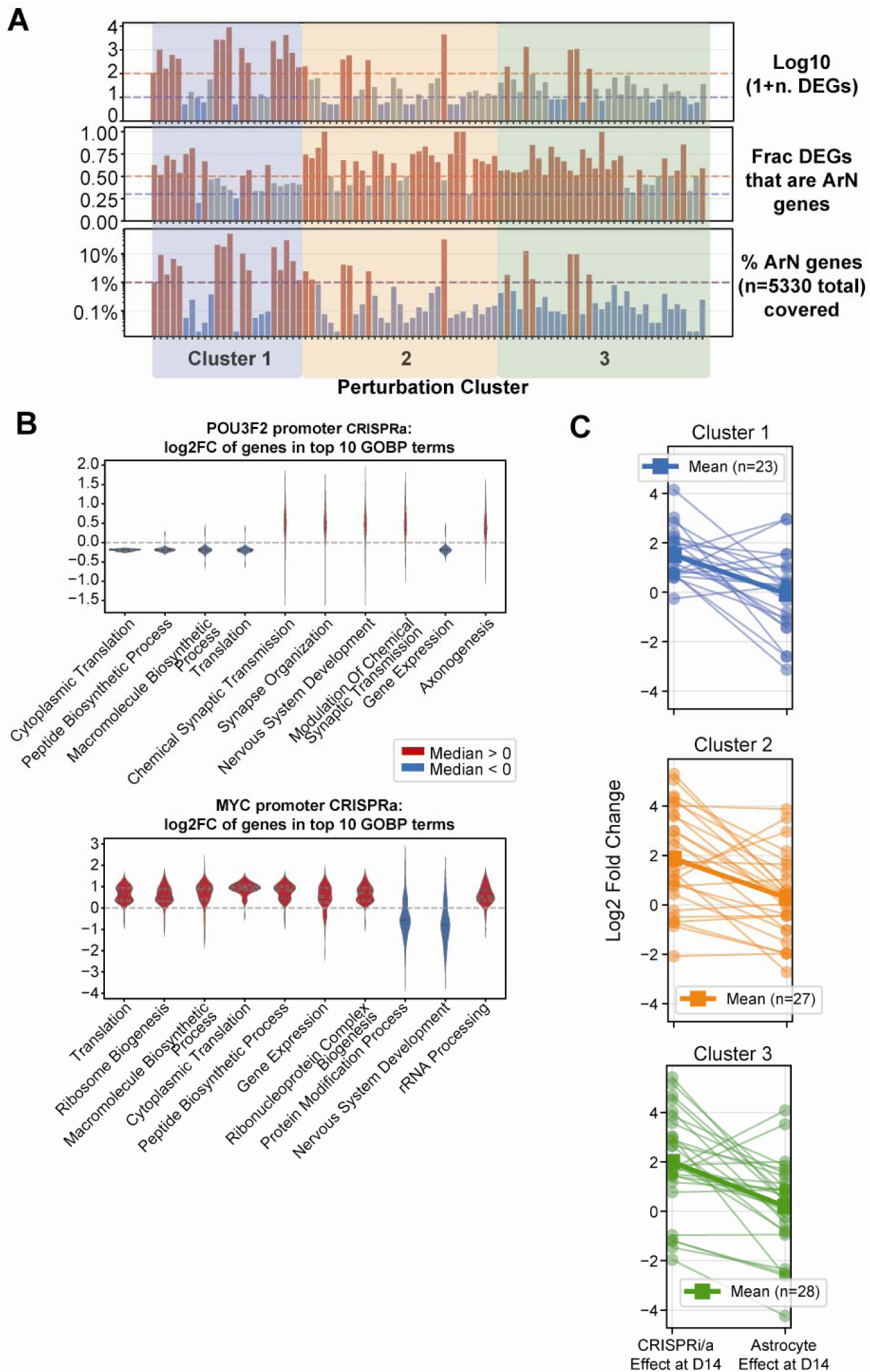
Additional metrics of the perturbation clusters identified in Fig. 4. (A) Three barplots showing perturbations in the same order as the columns of the heatmap in Fig. 4D showing three additional characteristics of the perturbations. **Top**, log10(number of global DEGs + 1). **Middle**, Fraction of global DEGs that are ArN genes. **Bottom**, % of ArN genes (n=5330, see (Fig. 1D)) covered by the global DEGs of this perturbation. (B) Violin plots showing the log2FC of global DEGs belonging to the top 10 GOBP terms of the described perturbation. Top, for POU3F2 promoter CRISPRa perturbation. Bottom, for MYC promoter CRISPRa perturbation. (C) Line charts showing, for each perturbation cluster, the comparison between the effects of CRISPR perturbations and astrocytes on target genes.

**Extended Data Figure 9:**
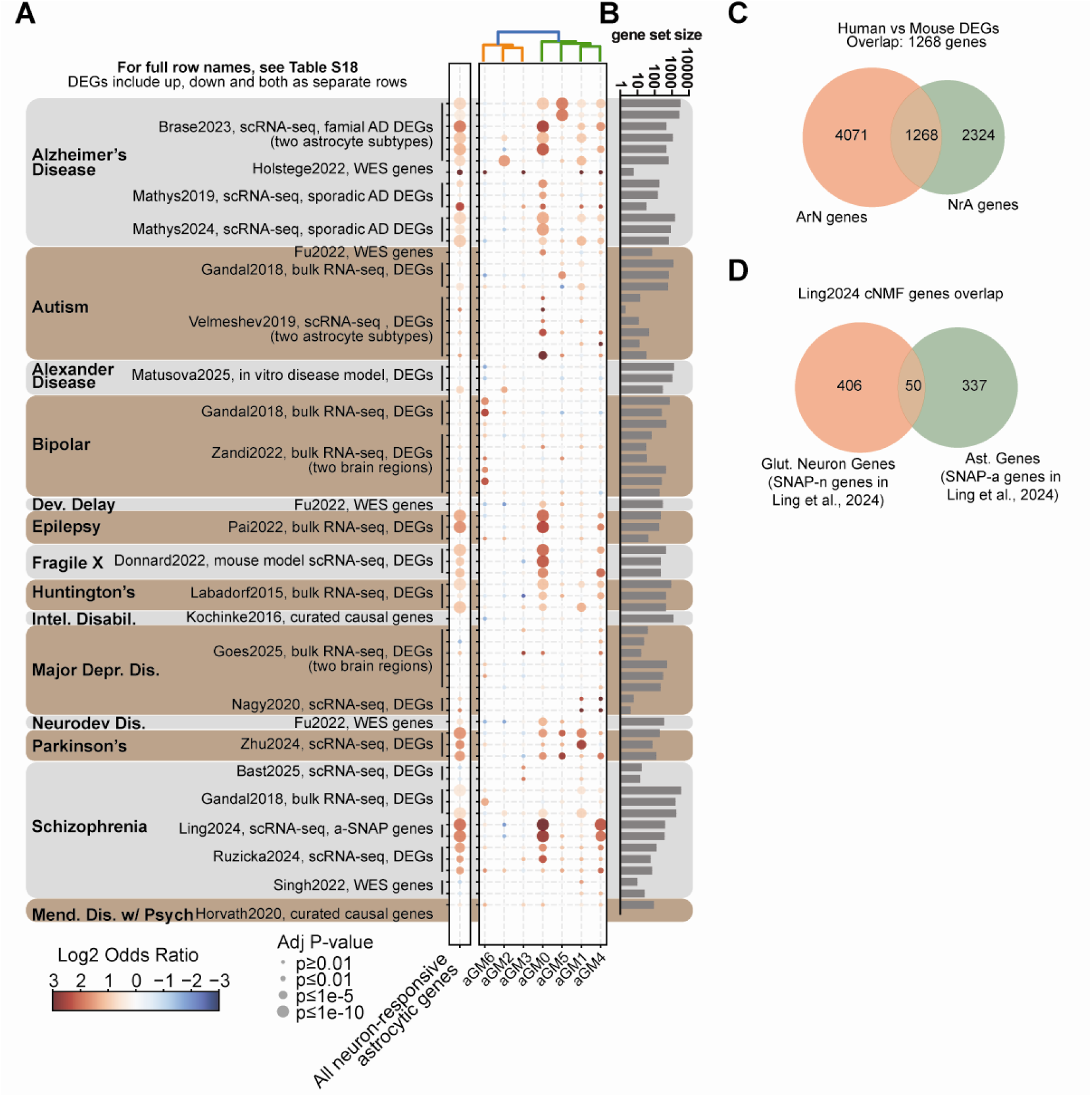
Neuron-responsive astrocytic gene programs are disrupted in diseases. (A) Dotplot showing the enrichment of NrA genes (union set, and individual nGMs, as described in **Extended Data** Fig. 2D) for various neurodevelopmental, neuropsychiatric, and neurodegenerative disease or disorder gene sets. Gene sets are labeled with publication (first author last name followed by year of publication), and name or category of the disease/disorder. A detailed list of the gene sets used can be found in **Table S18**. Abbreviations: WES, whole-exome sequencing-based gene set; bipolar, bipolar disorder; dev. delay, developmental delay; intel. disabil., intellectual disability; major depr. dis., major depressive disorder; neurodev. dis., neurodevelopmental disorders; mend. disease w/ psych, Mendelian disease with psychiatric symptoms. (B) Line chart showing the sizes of the disease gene sets. (C) Venn diagram showing the overlap between astrocyte-responsive neuron genes and neuron-responsive astrocyte genes. (D) Venn diagram showing the overlap between glutamatergic neuron- and astrocyte-contributed genes to the gene program found to be associated with schizophrenia and ageing in Ling et al., 2024.

**Extended Data Figure 10:**
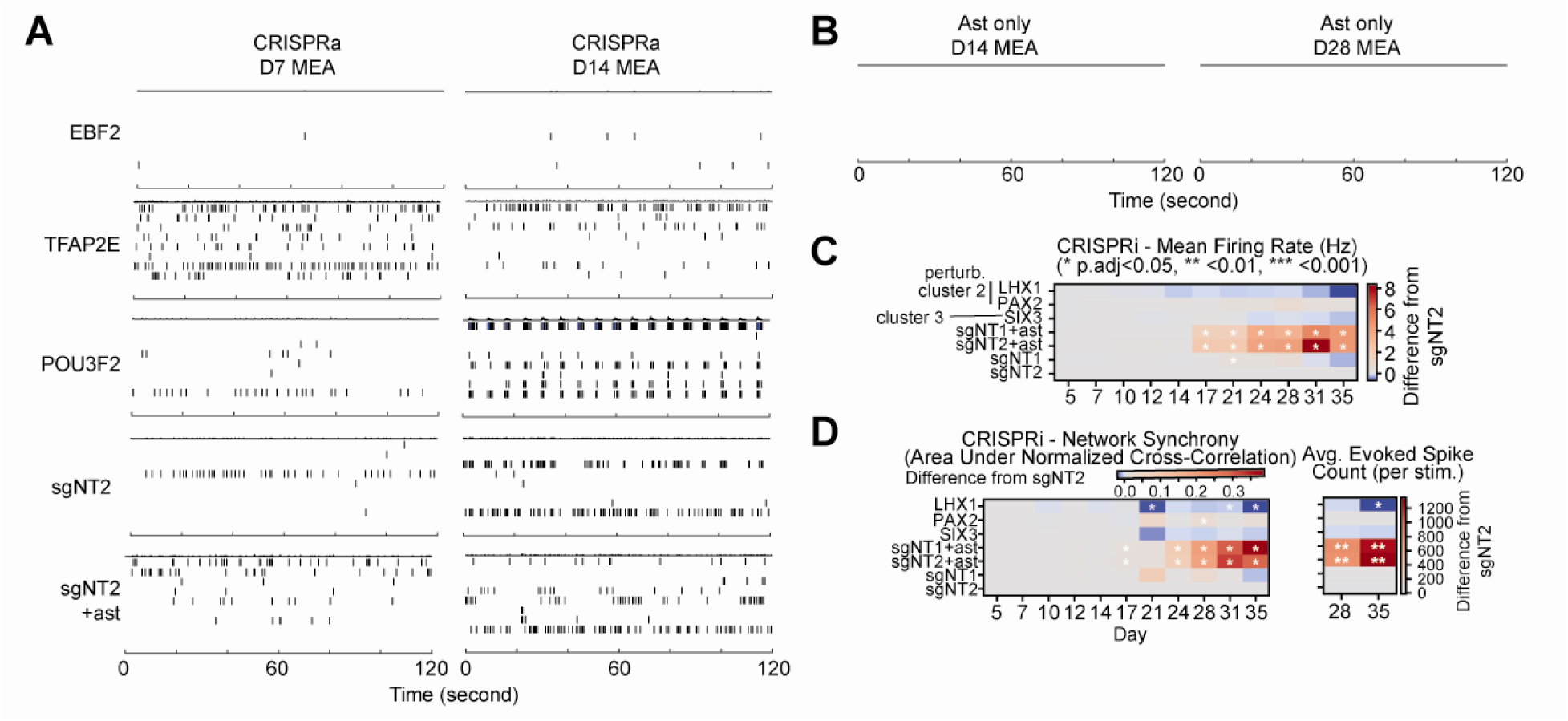
Additional electrophysiology data for ArN TF perturbations. (A) Raster plots showing the neurons’ firings on day 7 and day 14, across 8 electrodes of a representative well of 3 example CRISPRa perturbations, one negative control (sgNT2) and negative control co-cultured with astrocytes. (B) Raster plots showing the firings of astrocytes cultured in isolation on day 14 and day 24. (C) Heatmaps showing relative increase or decrease over negative control (sgNT2) of firing rate (Hz) of neurons with three individual CRISPRi fRE perturbations, negative control neurons, or neurons co-cultured with astrocytes. 8 replicates per condition were analyzed. (D) Heatmaps showing relative increase or decrease over negative control (sgNT2) of network synchrony (area under normalized cross-correlation) and average evoked spike count per stimulation of neurons with three individual CRISPRi fRE perturbations, negative control neurons, or neurons co-cultured with astrocytes. 8 replicates per condition were analyzed.

## Supplementary Table Legend

- **Supplementary Table 1**: Numbers of biosamples in bulk RNA-seq and ATAC-seq datasets
- **Supplementary Table 2**: A human gene table containing differential expression results of mono-vs co-cultured neurons from bulk RNA-seq
- **Supplementary Table 3**: Over-representation enrichment results of GOBP terms for bulk astrocyte-responsive neuronal gene programs (nGM0-8)
- **Supplementary Table 4**: Lists of marker genes examined to determine the purity of the isolated astrocytes (related to **Extended Data Fig. 2**)
- **Supplementary Table 5**: A mouse gene table containing differential expression results of mono-vs co-cultured astrocytes from bulk RNA-seq
- **Supplementary Table 6**: Over-representation enrichment results of GOBP terms for bulk neuron-responsive astrocytic gene programs (aGM0-6)
- **Supplementary Table 7**: Inferred receptor-ligand pairs mediating neuron-astrocyte interactions from a public single-cell brain atlas
- **Supplementary Table 8**: A table showing all (n=265,420) identified ATAC-seq peaks in mono- and co-cultured neurons, including the coordinates, annotation, and differential accessibility results
- **Supplementary Table 9**: Genomic Regions Enrichment of Annotations Tool (GREAT) enrichment results of GOBP terms for bulk astrocyte-responsive neuronal peak programs (nPM0-8)
- **Supplementary Table 10**: gkm-SVM performance metrics and motif weights for each astrocyte-responsive neuronal peak module (nPM0-8)
- **Supplementary Table 11**: Tables showing the 222 astrocyte-responsive neuronal TFs targeted in Perturb-seq and all pREs and additional promoter regions to those found in Horlbeck2016 that are targeted in Perturb-seq
- **Supplementary Table 12**: Tables of local analysis results for CRISPRi and CRISPRa pRE screens
- **Supplementary Table 13**: Tables of RT-qPCR sgRNA validation results of significant local pRE-gene links and the sgRNA sequences tested
- **Supplementary Table 14**: Tables showing fRE target genes that have been associated with diseases on ClinGen
- **Supplementary Table 15**: Result tables of global analysis (for effective promoter perturbations and fRE perturbations only) from all four Perturb-seq experiments
- **Supplementary Table 16**: A table showing the order of columns of the heatmap in **Fig. 4D**, along with the four metrics used to characterize them
- **Supplementary Table 17**: A table of datasets used in disease analysis
- **Supplementary Table 18**: Table showing the row names and descriptions of **Fig. 5A** and **Extended Data Fig. 9A**
- **Supplementary Table 19**: A table of fourteen perturbations selected for functional experiments, along with non-targeting controls used

## Methods

### Experimental methods

#### Human iPSC culture

Human iPSCs were maintained in complete mTeSR Plus medium (StemCell Tech, 100–0276). Cells were grown on tissue culture plastic pre-coated with Matrigel and kept in a humidified incubator at 37°C with 5% COL. For the coating, plates were incubated with Matrigel (Corning, 354230) diluted to 1 mg per 24 mL of DMEM/F12 (Gibco, 11320033) for a minimum of one hour at 37°C before use. For passaging, cells were treated with Accutase (StemCell Tech, 07920; Innovative Cell Technologies, AT104; Millipore, SCR005) and the medium was supplemented with 10 µM ROCK inhibitor (ROCKi; Y-27632, Stemcell Tech, 72304) for 16–28 hours.

### Engineering of monoclonal dCas9-p300 i3N-WTC11 iPSC line

We engineered a parental WTC-11 iPSC line that has an inducible mNgn2 knock-in cassette in the AAVS1 safe-harbor locus^15,16^ (i3N-WTC11 iPSCs; a gift from Dr. Yin Shen’s lab at UCSF) but expresses no CRISPR machinery. We used lentiviral integration to introduce the expression of dCas9-p300 (pLRB10) under the selection of puromycin. We derived monoclones by using CloneR (StemCell Tech, 05888) and manual colony picking. We verified dCas9-p300 genomic integration, mRNA expression, and gene activation at multiple regions using previously published sgRNAs^4,44^ and selected the best performing clone for Perturb-seq and downstream experiments.

### Differentiation of excitatory neurons

Excitatory neurons were derived from isogenic i3N-WTC11 iPSC cell lines as previously described ^15,16^. Briefly, i3N-WTC11 iPSCs were differentiated in two steps. First, they were pre-differentiated for 3 days with daily medium change on Matrigel-coated tissue culture plates (see **Methods** section ‘Human iPSC culture’) in knockout DMEM/F12 medium (Life Technologies, 12660-012) supplemented with 1× N-2 supplements (Life Technologies, 17502-048), 1× MEM NEAA (Life Technologies, 11140-050), 1 μg/mL mouse laminin (Life Technologies, 23017-015), 10 ng/mL BDNF (PeproTech, 450-02), 10 ng/mL NT3 (PeproTech, 450-03) and 2 μg/ml doxycycline hyclate (Sigma-Aldrich, D9891) (**Neuronal Pre-Differentiation Medium**). Second, the pre-differentiated cells were passed into post-mitotic maturation, at the beginning of which they were dissociated and re-seeded onto 10µg/mL poly-D-lysine-coated (PDL) (Sigma, P6407) tissue culture plates in medium containing equal parts DMEM/F12, HEPES (Life Technologies, 11330-032) and Neurobasal-A Medium (Life Technologies, 12349-015) supplemented with 0.5× B-27 supplements (Life Technologies, 17504-044), 0.5× N-2 supplements, 1× MEM NEAA, 0.5× GlutaMax (Life Technologies, 35050-061), 1 μg/mL mouse laminin, 10 ng/mL BDNF, 10 ng/mL NT3 and 2 μg/mL doxycycline hyclate (**Neuronal Maturation Medium**). Part of the medium is replaced every 7 days and doxycycline was omitted in the replacement medium.

### Isolation and culture of primary mouse cortical astrocytes

Similar to previously described^17^, primary cortical astrocytes were isolated from C57BL/6 mouse pups (P0.5-P1.5) and maintained in Astrocyte Growth Media (AGM) consisting of DMEM (Gibco, 11960-044) supplemented with 10% heat-inactivated fetal bovine serum (Gibco, 10437-028), 1% Penicillin-Streptomycin (Gibco, 15140148), 1% L-Glutamine (Gibco, 25030), 1% Sodium Pyruvate (Gibco, 11360), 5 mg/L Insulin (Sigma, I-6634), 5 mg/L N-Acetyl Cysteine (Sigma, A8199), and 5 mg/L Hydrocortisone (Sigma, H-0888) Cerebral cortices were dissected, minced (<1 mm³), and digested in DPBS (Gibco, 14287-080) containing Papain (Worthington, LK003176) and DNase I (Worthington, LS002007) for 45 minutes at 33°C with periodic agitation. The digestion was quenched using sequential low- and high-concentration ovomucoid inhibitor solutions containing BSA (Sigma, A8806) and Trypsin Inhibitor (Worthington, LS003086/LS003083), followed by trituration and filtration through a 40µm cell strainer [Corning, 352340]. Cells were seeded at high density (>30e6 cells/flask) into T75 flasks coated with 10 µg/mL Poly-D-Lysine (PDL) (Sigma, P6407). To remove non-astrocytic cells, cultures underwent vigorous shaking on day in vitro (DIV) 3, followed by treatment with Cytosine arabinoside (AraC) (Sigma, C1768) at a 1:2000 dilution on DIV5. On DIV7, purified astrocytes were detached using 0.05% Trypsin-EDTA (Gibco, 25300-054) and re-plated onto PDL-coated culture vessels, with or without neurons, for downstream assays.

### Co-culture of neurons and astrocytes

i3N-WTC11 iPSC-derived excitatory neurons (after entering maturation for one day) were co-cultured with mouse primary astrocytes (after being cultured in vitro for 7 or 8 days) at a 5:1 ratio on 10 µg/mL PDL-coated tissue culture plates after being dissociated with Accutase and 0.05% Trypsin-EDTA, respectively. For bulk RNA-seq and ATAC-seq, co-cultured cells were harvested after 2, 6, 13, and 27 days of co-culturing. For scRNA-seq, cells were harvested after 13 days of co-culturing.

### Lentivirus production

Lentivirus was produced in HEK293T cells cultured in Opti-MEM (Gibco, 31985088) supplemented with fetal bovine serum (Sigma-Aldrich, F2442), Sodium Pyruvate (Gibco, 11360070), MEM NEAA (Gibco, 11140050), and GlutaMAX (Gibco, 35050061). Cells were seeded at 7 x 10^6 cells per 10 cm dish and transfected 16 hours later using the Lipofectamine 3000 Transfection Reagent kit (Invitrogen, L3000008). A DNA mixture containing 9.75 µg psPAX2 (Addgene #12260), 3.25 µg pMD2.G (Addgene #12259), and 4.3 µg sgRNA plasmid pool was combined with P3000 reagent, mixed 1:1 with Lipofectamine 3000, and incubated for 15 minutes at room temperature. The complex was added to cells in 6 mL of medium (with 6mL of medium removed before transfection), which was replaced with 12 mL of fresh medium 6 hours post-transfection. Viral supernatants were harvested at 24 and 48 hours, pooled, and filtered through a 0.45 µm filter. Viral particles were concentrated using Lenti-X Concentrator (Takara, 631231) by incubating overnight at 4°C followed by centrifugation at 1,700 x g for 45 minutes. Pellets were resuspended in culture medium at 1/50th of the original volume.

### Lentiviral transduction of iPSCs and neurons

For the pRE Perturb-seq experiments, as well as the quantitative reverse transcription-PCR (qRT-PCR), immunofluorescence (IF), and multi-electrode array (MEA) assays, all of which were high-MOI, the following transduction protocol was used: iPSCs were dissociated into single cells using Accutase (Millipore, SCR005) for 6–7 minutes, centrifuged at 300 x g for 5 minutes, and resuspended in complete mTeSR Plus complete medium. Cells were counted using a Countess 3 Automated Cell Counter (Invitrogen) and seeded at a density of 1 x 10^6 cells per well in 6-well plates containing mTeSR supplemented with ROCK inhibitor. Lentiviral pools were immediately added to the wells at a high multiplicity of infection (MOI). Plates were gently agitated to ensure equal viral distribution and returned to the incubator. 24 hours later, iPSCs were pre-differentiated (see **Methods** section ‘Differentiation of excitatory neurons’).

For the low-MOI promoter Perturb-seq experiments, the following transduction protocol was used: Pre-differentiated neurons (Day 0) were dissociated using Accutase for 5 minutes, collected with PBS, pH7.4 (Gibco, 10010049), and centrifuged at 300 xg for 5 minutes. The cell pellet was resuspended in Neuronal Maturation Medium (see **Methods** section ‘Differentiation of excitatory neurons’) and counted using a Countess 3 Automated Cell Counter. Lentivirus was added directly to the bulk cell suspension to achieve a target transduction efficiency of 25%. The suspension was mixed by inversion and seeded onto PDL-coated 10 cm dishes at a density of 5 x 10^6 cells per dish.

### Cloning of individual sgRNAs

To clone the gRNAs into a lentivirus vector, we first constructed a lentiviral gRNA expression plasmid by combining a U6-gRNA cassette containing the gRNA-(F+E)-combined scaffold sequence with an EGFP-P2A-PAC cassette into a lentiviral expression backbone (Addgene #83925) using Gibson assembly (NEB, E2611).

Forward and reverse oligonucleotides representing the CRISPR sgRNA protospacer sequence were designed and synthesized (Integrated DNA Technologies or Genewiz) with 5’-CACCG-forward oligonucleotides-3’ and 5’-AAAC-reverse oligonucleotides-C-3’ architectures, respectively. Phosphorylation and annealing were performed by mixing 500 ng of each strand with T4 Polynucleotide Kinase (NEB, M0201) in T4 DNA Ligase Reaction Buffer (NEB, M0202), incubating at 37°C for 30 minutes, 95°C for 5 minutes, and ramping down to 25°C at 5°C/min. The annealed duplex was diluted 1:200 and cloned into Esp3I-digested (NEB, R0734) and gel-purified (Qiagen, 28706) lentiviral backbone (described above) via T4 DNA ligation (NEB, M0202). The reaction was incubated at 25C for one hour. The products were transformed into Stbl3 chemically competent E. coli and verified by Sanger sequencing (Genewiz). Stbl3 E. coli were created in house using the calcium chloride technique to achieve a transformation efficiency >1 x 107 cfu/μg. Cells were originally sourced from One Shot™ Stbl3™ Chemically Competent E. coli (Invitrogen, C737303) and renewed from the source periodically.

### Cloning of sgRNA libraries

The sgRNA oligonucleotide library was synthesized by Twist Bioscience using the following sequence format: ATATATCTTGTGGAAAGGACGAAACACCG-[19/20-bp protospacer sequence]-GTTTAAGAGCTATGCTGGAAACAGCATAG. The lentiviral backbone was linearized by digestion with Esp3I (NEB, R0734) and Calf Intestinal Alkaline Phosphatase (NEB, M0290) overnight at 37°C, followed by purification using SPRIselect beads (Beckman Coulter, B23318). The oligo pool was amplified via PCR using Kapa HiFi Hotstart ReadyMix (Kapa Biosystems, KK2602) for 10 cycles and gel-purified (Qiagen, 28706). The purified insert and backbone were assembled using Gibson Assembly Master Mix (NEB, E2611) and electroporated into Endura Electrocompetent Cells (Lucigen, 60242-2). Transformed cells were plated onto large bioassay plates and grown at 37°C for 16 hours. Colonies were harvested, and plasmid DNA was extracted using the Qiagen Plasmid Midi Kit (Qiagen, 12143). Library distribution and diversity were subsequently verified by Next Generation Sequencing.

### Bulk RNA-seq assay

Neuron and astrocyte monocultures and neuron-astrocyte co-cultures were collected for RNA extraction after neurons entered the maturation phase for 3, 7, 14, and 28 days (i.e., 2, 6, 13, and 27 days after co-culturing, respectively). Astrocyte monoculture was incubated in the identical medium as neuron mono- and co-cultures. RNA extraction followed the manufacturer’s recommended protocol for silicaLcolumn purification (RNeasy Mini Kit, Qiagen 74104), with some experiments incorporating an onLcolumn DNase digestion step to remove residual genomic DNA (RNaseLfree DNase Set, Qiagen 79254). RNA was eluted in RNaseLfree water in small volumes (typically 30 µL, occasionally as two sequential elutions), and concentrations were measured using a fluorometric assay (Qubit RNA Broad Range Assay Kit, Invitrogen Q10211) on Invitrogen Qubit 4 Fluorometer.

RNA samples were then shipped to Genewiz for RNA-seq library prep (Library preparation, Illumina, RNA with rRNA depletion) and sequencing (Illumina, 2×150bp, ∼350M PE reads) services, with targeted sequencing depth for the mono-cultured neurons and astrocytes samples at 30 million paired-end reads per sample, and for the co-cultured neurons and astrocytes samples at 60 million per sample.

### Bulk ATAC-seq assay

Neuron and astrocyte monocultures and neuron-astrocyte co-cultures were dissociated (for details, see **Methods** section ‘Dissociation of neuronal cultures for flow cytometry or single-cell RNA-seq’) and viably frozen in 90% KnockOut Serum Replacement (Gibco, 10828010) + 10% dimethyl sulfoxide (Sigma-Aldrich, D8418) after neurons entered the maturation phase for 3, 7, 14, and 28 days (i.e., 2, 6, 13, and 27 days after co-culturing, respectively). Approximately 100,000-500,000 viably frozen cells were used for ATAC-seq library preparation according to the Omni-ATAC protocol ^92^ with modifications. Frozen vials were thawed at 37C. Cells were washed with warm growth media then washed with 1X PBS. Cell pellets were suspended in 1mL of ATAC-seq RSB containing 0.1% Tween and transferred to a dounce homogenizer. Cells were lysed with 10 stokes (tight pestle) and centrifuged for 10 min at 500 rcf. Nuclei pellets were suspended in 27μl 2X TD buffer and 2ul were used for counting using hemocytometer. 50,000 nuclei were added to the transposition reaction mix (2.5 μl transposase, 16.5 μl PBS, 0.5 μl 1% digitonin, 0.5 μl 10% Tween-20, 5 μl water, and 2X TD buffer to 50ul) and incubated at 37 °C for 30 min in a thermomixer with shaking at 1,000 rpm. Reactions were cleaned up with Zymo DNA Clean and Concentrator-5 kit. ATAC-seq libraries were double indexed with Nextera PCR Primers and amplified with 9 to 12 cycles of PCR. Amplified DNA fragments were purified with 1:1 ratio of Agencourt AMPure XP (Beckman Coulter). Libraries were quantified by Qubit and size distribution was inspected by Tapestation (Agilent High Sensitivity DNA, Agilent Technologies).

### Dissociation of neuronal cultures for flow cytometry or single-cell RNA-seq

Neuronal cultures were dissociated using a papain-based protocol. A digestion solution was prepared by combining Papain (Worthington, LK003178) in Accutase (Millipore, SCR005) to achieve final concentrations of 20 units/mL and supplementing it with 10 µM ROCK inhibitor. The solution was sterile-filtered and pre-activated at 37°C for 10 minutes. Cultures were washed with DPBS (Gibco, 14190144) and incubated with the papain solution for 40 minutes at 37°C. The reaction was quenched by transferring the sheet(s) of neurons, which can be easily detached from the tissue culture vessel mechanically at this point, into 5mL Neuronal Maturation Medium (see **Methods** section ‘Differentiation of excitatory neurons’) supplemented with 250 units/mL DNase I (Worthington, LK003172) and 10 µM ROCK inhibitor. Cells were triturated gently by pipetting, and centrifuged at 300 x g for 5 minutes. For flow cytometry, the pellet was resuspended in DPBS (Sigma-Aldrich, D8537) + 1% bovine serum albumin (Sigma, A0281) and the neurons were then subject to flow cytometry. For scRNA-seq, the cells were washed two more times in Neuronal Maturation Medium + 1% bovine serum albumin (Sigma, A0281) and filtered through a 30 µm Pre-Separation Filter (Miltenyi Biotec, 130-041-407) prior to loading.

### Single-cell RNA-seq of neuron mono-culture or neuron-astrocyte co-culture

Neuron mono-culture or co-culture were subject to the protocol described under **Methods** section ‘Dissociation of neuronal cultures for flow cytometry or single-cell RNA-seq’ on day 14 of maturation phase. Single-cell suspensions were adjusted based on post-filtration counts and loaded onto a 10x Genomics Chromium Next GEM 5′ HT v2 Chip N with 5’ CRISPR kit to allow sgRNA capture, following the manufacturer’s protocol (10X Genomics, PN-1000375, PN-1000451). Loading volumes were calculated by assuming effective concentrations ∼1.2-fold lower than measured, to target ∼20,000 cell recovery per lane with ∼20% overloading (e.g., treating a cell suspension concentrated at 1000 cells/µL as 800 cells/µL). Single-cell gene expression and sgRNA expression libraries were generated using the 10x Genomics Next GEM 5’ HT v2 and 5’ CRISPR kits following the manufacturer’s protocols (10X Genomics, User Guide CG000512 Rev B, PN-1000374, PN-1000375, PN-1000451, PN-1000215). Final libraries were quality checked based on concentration (Qubit 4 Fluorometer; Qubit dsDNA High Sensitivity Assay Kit, Invitrogen Q32854) and fragment-size distribution (Agilent TapeStation 2200 or 4200; DNA D1000 and D5000 assays, Agilent 5067-5582, 5067-5583, 5067-5588 5067-5589) and were used for downstream Illumina Nextseq 2000, NovaSeq 6000 or NovaSeq X Plus sequencing to target depths of ∼5 x 10^4 and ∼5 x 10^3 reads/cell for the gene expression and sgRNA expression libraries, respectively.

### Design of pRE Perturb-seq screens

We first identified all astrocyte-responsive pREs (peaks with FDR < 0.01 shown in **Fig. 2B**) located within ±100kb from any TSS of the 222 astrocyte-responsive TF genes (shown in **Extended Data Fig. 5A**, **Table S11**; defined in^93,94^). To avoid targeting promoters, we excluded the strongest ATAC peak within ±5kb from the TSSs of each TF gene from the screens. There are in total 735 pREs that fit these criteria. They were further split into two non-overlapping sets based on their chromatin accessibility’s mean direction of change across D3, D7, and D14 (**Fig. 2B**). Those that lose accessibility upon co-culture with astrocytes were targeted with CRISPRi (dCas9-KRAB), and those that gain accessibility were targeted with CRISPRa (dCas9-p300). This resulted in 342 CRISPRi-targeted pREs, and 393 CRISPRa-targeted pREs (**Extended Data Fig. 6B**).

We designed two sgRNA libraries that target the CRISPRi- and CRISPRa-targeted sets of pREs, respectively, with guidescan2 (specificity score >0.1 for ‘no-alt’ version of hg38) ^95^, for an average of nearly 10 sgRNAs per pRE. To strike a balance between spreading out sgRNAs targeting the same region to minimize overlapping sgRNAs which could potentially result in similar off-targeting profiles, and concentrating sgRNAs around ATAC-seq peak summits to maximize efficacy ^96^, we selected 1, 2, 2, 2, 2, 1 gRNA, respectively, for the 0-20%, 20-40%, 40-

50%, 50-60%, 60-80% and 80-100% portions from 5’ to 3’ across each target pRE. For each portion, sgRNAs with the highest guidescan2 efficiency score were selected. We further included 5% of previously published bona fide non-targeting sgRNAs as negative controls ^96^ and a few over-represented positive control sgRNAs that target promoter regions of additional genes (TFRC, UBQLN2, GRN, and CDH2 for CRISPRi ^97^; GRIN2A, CXCR4, COL4A1, and MYOD1 for CRISPRa ^44^), and in the case of dCas9-p300 library, also a distal regulatory region that was used in the validation of the p300 cell line (**Extended Data Fig. 5D**) ^4^. This resulted in 3,344 and 4,130 sgRNAs for the CRISPRi and CRISPRa libraries, respectively.

### pRE Perturb-seq experiments

i3N-WTC11 iPSCs constitutively expressing dCas9-KRAB and dCas9-p300, respectively, were transduced at high MOI with lentiviral sgRNA libraries that target pREs that lose or gain accessibility upon astrocyte co-culture. Cells were pre-differentiated 24 hours later. For details, see **Methods** section ‘Lentiviral transduction of iPSCs and neurons’. On day 14 of maturation, neurons were dissociated and subject to loading for scRNA-seq and single-cell gene expression and sgRNA expression library prep according to **Methods** section ‘Single-cell RNA-seq of neuron mono-culture or neuron-astrocyte co-culture’. In addition, the gene expression libraries were subjected to target enrichment using a hybridization-based method (Twist Bioscience, PN 101001, 104445, 100856, and 101262) with a panel of DNA probes targeting the 5’ UTR and first ∼400bp of the coding sequence of all transcripts (Gencode v41) of the 222 TF genes of interest (**Table S11**) along with 4 positive control genes (TFRC, UBQLN2, GRN, and CDH2) as well as 7 housekeeping genes (DPM1, KRIT1, BAD, ARF5, POLDIP2, HCCS, TSR3). Target enrichment was performed according to the Standard Hybridization v2 Protocol (Twist Bioscience, DOC-001273 REV 3.0). Briefly, pooled libraries were concentrated by vacuum centrifugation and resuspended in a mixture containing Twist Universal Blockers and Twist Blockers. The library pool was hybridized with biotinylated probes in Twist Hybridization Mix at 70°C for 16 hours. Target-probe complexes were captured using Streptavidin Binding Beads and washed with pre-heated Twist Wash Buffers. The enriched libraries were amplified using the 10X Genomics Amp Mix (10X Genomics, PN 2000047) and purified using Twist Purification Beads. Finally, the target-enriched gene expression libraries were sequenced using Illumina Nextseq 2000, NovaSeq 6000 or NovaSeq X Plus sequencing to target depth of ∼5 x 10^3 reads/cell.

### Design of promoter Perturb-seq experiments

The 222 astrocyte-responsive TF genes (shown in **Extended Data Fig. 5A**, **Table S11**) were split into two non-overlapping sets based on their mRNA level’s mean direction of change across D3, D7, D14 and D28 (**Fig. 1D**). Those that lose mRNA expression upon co-culture with astrocytes were targeted with CRISPRi (dCas9-KRAB), and those that gain mRNA expression were targeted with CRISPRa (dCas9-VPH ^48^). This resulted in 145 CRISPRi target TF genes, and 77 CRISPRa target TF genes.

We designed two sgRNA libraries to target CRISPRi and CRISPRa to the promoters of these sets of TF genes, respectively, by subsetting previously published human genome-wide gene promoter-targeting libraries (hCRISPRi-v2.1 and hCRISPRa-v2) ^44^. One TF gene in each set (ZNF204P in CRISPRi set and FOXD4 in CRISPRa set) was not found in the published libraries and was thus removed from the experiment.. Some TFs have multiple promoters (25 of 144 in CRISPRi set; 13 of 76 in CRISPRa set), and all were included in the experiments. 10 sgRNAs per promoter were included. In addition, 13 and 8 unannotated putative promoters were respectively identified of the CRISPRi and CRISPRa sets of TF genes by manually inspecting our bulk RNA-seq and ATAC-seq data (by two criteria: **a,** an ATAC-seq peak is present, and **b,** RNA transcription initiates from the ATAC-seq peak). For these additional targets, we designed 10 sgRNAs per promoter based on the ATAC-seq peak call with guidescan2 (specificity score >0.2 for ‘no-alt’ version of hg38; the 10 sgRNAs with the highest efficiency scores were selected) ^95^. We further included 5% of previously published bona fide non-targeting sgRNAs as negative controls ^96^ and four positive control promoters ( of the genes TFRC, UBQLN2, GRN, and CDH2 for both CRISPRi and CRISPRa; 10 sgRNAs per promoter were taken from the hCRISPRi-v2.1 and hCRISPRa-v2 libraries, respectively). This resulted in 1,968 and 1,063 sgRNAs for the CRISPRi and CRISPRa libraries, respectively (**Extended Data Fig. 7B**).

### Promoter Perturb-seq experiments

Pre-differentiated i3N-WTC11 iPSCs constitutively expressing dCas9-KRAB or inducibly expressing dCas9-VPH ^48^, respectively, were transduced at low MOI with lentiviral sgRNA libraries that target TF gene promoters that lose or gain mRNA expression upon astrocyte co-culture. Cells were dissociated and plated for maturation as lentivirus was introduced. For the dCas9-VPH experiment, 20µM trimethoprim (TMP) (Sigma, 92131) was added at the same time as the lentivirus, as well as at medium change, to induce the stabilization of the dCas9-VPH protein. For more details of the transduction method, see **Methods** section ‘Lentiviral transduction of iPSCs and neurons’. On day 14 of maturation, neurons were dissociated (for details, see **Methods** section ‘Single-cell RNA-seq of neuron mono-culture or neuron-astrocyte co-culture’) and subject to fluorescence-activated cell sorting (FACS) on a Sony Cell Sorter SH800Z to separate out and collect transduced (EGFP+, EGFP co-expressed with sgRNA) neurons. The EGFP+ neurons were then subject to loading for scRNA-seq and single-cell gene expression and sgRNA expression library prep according to **Methods** section ‘Single-cell RNA-seq of neuron mono-culture or neuron-astrocyte co-culture’.

### RNA extraction for qRT-PCR

For all qRT-PCR assays, i3N-WTC11 iPSCs expressing the appropriate dCas9-effector protein were transduced at high MOI with single lentiviruses expressing a sgRNA, pre-differentiated, and matured. For each sgRNA, 3 biological replicates were set up. Neurons were harvested at 14 days of maturation for RNA extraction, with the exception of the dCas9-p300 cell line’s validation experiment (**Extended Data Fig. 5D**), which was harvested 7 days after maturation. RNA extraction was performed with the Norgen Total RNA Purification Plus Micro Kit (Norgen Biotek 48500). After medium removal, cells were lysed with Buffer RL on plate. Cells lysates were optionally stored at −80C before further processing. Cell lysates were then processed according to the manufacturer’s instructions.

### RT-qPCR

RNA samples were reverse transcribed (RT) with the SuperScript VILO cDNA Synthesis Kit (Invitrogen, 11754250) following the manufacturer’s instructions. 100 - 500ng of RNA was used as input for each 20µL reaction. The RT products were diluted in molecular biology-grade water (Corning, 46-000-CM) for 1:10. The diluted products were then subject to quantitative PCR (qPCR) reactions.

qPCR were set up with PerfeCTa SYBR Green FastMix (QuantaBio, 95072) for primer-based reactions (for genes GRIN2A and OLIG3) or PerfeCTa FastMix II (QuantaBio, 95118) for TaqMan Gene Expression Assay (Applied Biosystems, 4448892 or 4453320)-based reactions (for all other genes). The gene TBP was included in both types of assays for each RNA sample as the housekeeping control gene. Reactions were loaded on a 96-well plate (Thermo Scientific, AB0700W) and run on a CFX96 Deep Well Real-Time System mounted on a C1000 Touch Thermal Cycler (Bio-Rad). Ct values exported from the Bio-Rad CFX Maestro software for analysis.

### Immunofluorescence and microscopy

For staining of neurons with CRISPRi/a perturbations, i3N-WTC11 iPSC cell lines stably expressing dCas9-KRAB for CRISPRi or inducibly expressing dCas9-VPH by TMP for CRISPRa were maintained in mTeSR plus medium on Matrigel-coated tissue culture plates under standard conditions (37 °C, 5% COL). iPSCs were transduced with individual lentiviruses expressing sgRNAs for the intended CRISPRi or CRISPRa perturbations, along with two non-targeting sgRNAs (**Table S19**) were transduced at high multiplicity of infection using virus-containing media prepared by diluting thawed viral supernatants into mTeSR (+ROCKi) to the desired working concentration. After overnight exposure, cells were washed and transitioned into pre-differentiation medium for subsequent differentiation. At the switch to Neuronal Maturation Medium, cells were plated on µ-Slide 8 Well chambered coverslips (Ibidi, 80804 or 80826) that were pre-coated with PDL only or PDL followed by 20µg/mL laminin (Gibco, 23017015) for 1.25e5 cells per well in 250µL of medium. TMP was included in the medium at plating and medium change for dCas9-VPH cells to stabilize CRISPRa components. For cells transduced with either of the two non-targeting sgRNAs, co-culture with astrocytes were also set up, with 2.5e4 astrocytes co-plated with neurons in an additional 50µL of medium.

After switching to Neuronal Maturation Medium for 7 or 14 days, cells on ibidi chamber slides were rinsed in DPBS and fixed in warmed 4% paraformaldehyde (PFA) (Electron Microscopy Sciences, 15710) for 15 minutes at room temperature. Following three washes with PBS, cells were blocked and permeabilized for 30 minutes at room temperature in a solution containing 50% Normal Goat Serum (NGS) (Abcam, ab7481), 0.2% Triton X-100 (Sigma-Aldrich, X100PC), and 50% base buffer (DPBS with 1% BSA). Primary antibodies (anti-GFP, Sigma-Aldrich AB16901; anti-βIII-tubulin, Biologend 802001; anti-SYN1, Sigma AB1543; anti-MAP2, Sigma-Aldrich Sigma AB1543; anti-GFAP, Proteintech 16825-1-AP) were diluted in a buffer containing 10% NGS and 90% base buffer and applied overnight at 4°C. After washing, cells were incubated with Alexa Fluor-conjugated secondary antibodies (Invitrogen) diluted in the same buffer for 1 hour at room temperature protected from light. Nuclei were stained with DAPI (Invitrogen, 62248) (1:1,000) before covering each well with 150µL Vectashield (VWR, H-1000-10). Images were acquired using a Zeiss Axio Observer microscope with a 20x objective. CZI files were exported through the Zeiss Zen Pro software for downstream processing.

### Multi-electrode array (MEA)

iPSC cells were maintained, transduced, and differentiated the same way as described in Methods section, ‘Immunofluorescence and microscopy’. At the switch to Neuronal Maturation Medium, cells were plated on MEA 96-well plate (Axion Biosystems M768-tMEA-96W). To prepare the MEA plate, all wells that are to be used were first coated with 150µL/well fetal bovine serum for 60 minutes at 37C. Then, FBS was aspirated and the wells were washed three times with PBS. The wells were then coated with PDL for 60 minutes at 37C. Then, PDL was aspirated and the wells were washed three times with PBS. Next, the plate was left open in the biosafety cabinet to air dry for 60 minutes before plating cells. 5µL of 1e7/mL (5e4 total) pre-differentiated neurons resuspended in Neuronal Maturation Medium (with Dox, and with TMP for dCas9-VPH cells) were plated at the center of each well covering the area of the well with the electrodes. For neuron-astrocyte co-cultures, an additional 1µL of 1e7/mL (1e4 total) astrocytes were co-plated with the neurons. Plate was incubated at 37C for 45 minutes for the cells to attach, and 150µL/well more warm Neuronal Maturation Medium was gently added at the end of the incubation. MEA recording was taken 2-3 times a week for viability and spontaneous activity on Axion Maestro Pro. Neurons were fed as described in **Methods** section ‘Differentiation of excitatory neurons’, with TMP included in the medium for the dCas9-VPH cells at the day-7 feeding, but not in the later feedings. Recording was taken for up to five weeks with the AxIS Navigator 3.10.1 software. On day 28 and 35, after recording spontaneous activity, electrical stimulation (using a Neural Stimulation block in the Stimulation Studio in AxIS Navigator that sends biphasic rectangular voltage pulses) at 1V for 1ms for 4 of 8 electrodes per well once every 20 seconds was applied to all wells for 10 minutes and the response to the stimulus was recorded. Spike files were generated automatically by the software at the end of each recording and were used for downstream analysis.

## Computational and statistical methods

### Analysis of bulk RNA-seq

#### Preprocessing of bulk RNA-seq

Raw reads were first mapped to both human reference GRCh38 (gene annotation Gencode v41) and mouse reference GRCm38 (gene annotation Gencode m25) genomes using STAR (v2.5.3) ^98^. The resulting BAM files were determined to be either human or mouse using XenofilteR ^27^. The species-confident reads were then converted to fastq files by bedtools (v2.30.0) ^99^. Reads assigned to be mouse are assumed to come from astrocytes; reads assigned to be human are assumed to come from neurons. The appropriate fastq files for neuron monoculture, astrocyte monoculture, and neuron-astrocyte co-culture were then processed with the ENCODE RNA-seq pipeline (https://github.com/ENCODE-DCC/rna-seq-pipeline) ^28^.

### Differential expression analysis and Principal Component Analysis of bulk RNA-seq by DESeq2

Differential gene expression analysis was performed using the DESeq2 package in R ^29^. Raw ‘expected count’ from the ENCODE RNA-seq pipeline were rounded to integers prior to analysis. A likelihood ratio test (LRT) was conducted using a full design model accounting for days of maturation, culture condition (mono-vs co-culture), experiment batch, and the interaction between day and condition (∼ day + culture_condition + batch + day:culture_condition) compared against a reduced model (∼ day + experiment). Low-abundance genes with a mean expression (baseMean) less than 3 were excluded from the results. Significant differentially expressed genes were identified based on a.adj < 0.01.

Normalized read counts were extracted from the DESeq2 object using the counts() function with normalized=TRUE for downstream analysis. For dimensionality reduction and quality assessment, variance-stabilizing transformation (vst) was applied to the raw count data, and Principal Component Analysis was performed using DESeq2’s plotPCA() function separately for each batch variable (day, culture_condition, batch) to visualize sample clustering.

### Pairwise correlation between effect sizes at different time points of co-culture-responsive genes

Correlation analysis was restricted to genes meeting statistical (padj < 0.01) and abundance thresholds (baseMean ≥ 3), thousands of genes. Samples were grouped by day of differentiation and culture condition, and expression values (DESeq2-normalized read counts) were averaged across biological replicates within each group. Culture-condition ratios (co-cultured neurons/mono-cultured neurons, or co-cultured astrocytes/mono-cultured astrocytes) were calculated for each timepoint, then converted to log2 scale. Pearson correlation coefficients were computed among all pairs of time points for these differential genes.

### Unsupervised clustering of differentially expressed genes

Differentially expressed genes (padjL<L0.01, count-filtered) by zLscored log1p-normalized RNA counts using Scanpy ^100^. An AnnData matrix was built with genes as observations and samples as variables, followed by PCA (20 components), kLnearest neighbors on the PCA space, Louvain community detection (resolutionL 0.75), PAGA graph abstraction, and UMAP for visualization. Genes were ordered by Louvain cluster membership to generate heatmaps in Seaborn (clustermap) with cluster-specific row colors.

### Gene ontology over-representation analysis for gene clusters (modules)

We performed over-representation analysis on Louvain-derived gene modules from **Methods** section ‘Unsupervised clustering of differentially expressed genes’. Background genes were defined as all expressed genes with baseMean ≥ the minimum among padj□<□0.01 DEGs. For each cluster’s gene list, we ran Enrichr via gseapy ^101^ against GO Biological Process 2025, SynGO 2024, WikiPathways 2024 Human, Reactome Pathways 2024, and KEGG 2021 Human, supplying the background list. Significant over-enrichment was identified with adjusted p < 0.05.

### Comparison with previous publications

Astrocyte-responsive neuron genes were overlapped with neuron genes found to respond to glia in two other studies ^19,20^. First, a ‘gene universe’ was defined based on the DESeq2 baseMean >3 filter as described in **Methods** section ‘Differential expression analysis of bulk RNA-seq’. Only genes from the other publications that intersect the ‘gene universe’ were kept. Then, pairwise and three-way overlaps were identified, and enrichment over chance was calculated by hypergeometric test. Similarly, neuron-responsive astrocyte genes were overlapped with astrocyte genes found to respond to neurons ^33^ or with astrocyte maturity genes^32^.

### Analysis of bulk ATAC-seq

#### Preprocessing of bulk ATAC-seq

Similar to RNA-seq, raw reads were mapped to both human reference GRCh38 and mouse reference GRCm38 using BWA (v0.7.17) ^102^, assigned to either using XenofilteR, converted back to fastq using bedtools, and processed with the ENCODE ATAC-seq pipeline (https://github.com/ENCODE-DCC/atac-seq-pipeline) that is modified to output peak calls with FDR <0.01 instead of the default IDR method. A reproducible union peak set was called across all samples with MSPC ^103^, requiring a peak to be called in at least two samples to be kept for downstream analysis. This resulted in a set of 265,420 peaks. Finally, featureCounts ^104^ was used to count the numbers of reads overlapping the peaks for each sample. Because D28 samples had low data quality, they were removed from downstream analysis.

Astrocyte data did not pass quality control and were discarded, which we reasoned not to be a major inadequacy, given our focus on neurons in this report for their relevance to human biology and the availability of epigenome editing tools^16,48^ that enable the perturbation experiments described later.

### Differential accessibility analysis and Principal Component Analysis of bulk ATAC-seq by DESeq2

Similar to RNA-seq, differential chromatin accessibility analysis was performed using the DESeq2 package in R using peak counts from featureCounts. The same formulae and parameters were used as RNA-seq. Normalized read counts were similarly extracted and variance-stabilizing transformation (vst) was applied to the raw count data before Principal Component Analysis was performed.

### Pairwise correlation between effect sizes at different time points of co-culture-responsive ATAC-seq peaks

The same method as ‘Pairwise correlation between effect sizes at different time points of co-culture-responsive genes’ was used.

### Unsupervised clustering of differentially accessible ATAC-seq peaks

Differentially accessible peaks (padjL<L0.01, count-filtered) by zLscored log1p-normalized read counts using Scanpy. An AnnData matrix was built with genes as observations and samples as variables, followed by PCA (20 components), kLnearest neighbors on the PCA space, Louvain community detection (resolutionL 0.75), PAGA graph abstraction, and UMAP for visualization. Peaks were ordered by Louvain cluster membership to generate heatmaps in Seaborn (clustermap) with cluster-specific row colors.

### Gene Ontology GREAT analysis for ATAC-seq peak clusters (modules)

Each peak module was functionally annotated using Genomic Regions Enrichment of Annotations Tool (GREAT) analysis with the union peak set used as background ^40,105^. A bed file representing genomic coordinates of peaks in each module was created and used as input for GREAT analysis against the gene set library GO Biological Process. Significant GO BP terms were filtered to adjusted p < 0.05 and a composite significance score was calculated per term as -log10(adjusted p) X |log2(fold enrichment)| X log2(gene set size), then ranked to prioritize high-impact pathways for downstream interpretation.

### Nearest-gene association analyses of astrocyte-responsive neuron ATAC-seq peaks

For association between all astrocyte-responsive neuron genes and ATAC-seq peaks, the following was done: first, each astrocyte-responsive neuron ATAC-seq peak was associated with its nearest gene by ChIPseeker ^106^. Next, the mean response (i.e., mean log2fc(co-cultured neuron/mono-cultured neuron)) of each astrocyte-responsive neuron ATAC-seq peak and gene across D3, D7, and D14 was determined and categorized into either up (mean log2fc>0) or down (mean log2fc<0). Finally, the average numbers of up and down peaks for up and down genes were calculated.

For association between gene and ATAC-seq peak modules, the distance-based gene-peak associations were preserved. Louvain-derived gene modules (detailed in Methods section ‘Unsupervised clustering of differentially expressed genes’) and their associated astrocyte-responsive ATAC-seq peaks were mapped to Louvain-derived peak modules (detailed in Methods section ‘Unsupervised clustering of differentially accessible ATAC-seq peaks’) based on gene-peak associations. An observed count matrix was constructed with gene modules as rows and peak modules as columns, tabulating the number of peaks from each peak module associated with genes in each gene module. Expected counts were calculated under the null hypothesis of random association using the hypergeometric distribution: for each gene-peak module pair, expected = (number of peak draws per gene module) × (peak module size / total number of peaks). Observed/expected ratios were log2-transformed to create fold-enrichment matrices. Statistical significance of observed vs. expected associations was assessed using chi-squared tests of independence on 2×2 contingency tables for each gene-peak module pair. P-values were adjusted for multiple testing using the Benjamini-Hochberg method within each gene cluster.

### gkm-SVM of ATAC-seq peak modules

gkm-SVM was trained on clusters vs negative genomic sequences as described in ^42^ using the gkm-SVM R package ^43^ with default parameters. Motifs were mapped to TFs using gkm-PWM [ref: Shigaki D, Yang Y, Eng NWL, and Beer MA. Systematic identification of cell-specific transcription factor activity and binding site mapping in 1644 human ENCODE chromatin accessibility datasets. In revision in Nature Genetics, 2026] (http://beerlab.org/gkmpwm/motif_gene_list).

### scRNA-seq analysis of mono- and co-cultured neurons

#### Data preprocessing and quality filtering of cells

Single-cell transcriptomics data were processed using a multi-step quality control and preprocessing pipeline implemented in Python (v3.10.12) with Scanpy (v1.9.6), AnnData (v0.10.3), and related libraries (numpy v1.26.4, pandas v2.1.4, scipy v1.11.4). Raw gene expression matrices were generated by Cell Ranger (10X Genomics, v7.2.0) using the human/mouse combined reference (refdata-gex-GRCh38_and_GRCm39-2024-A) from fastq files generated by Illumina NextSeq2000, NovaSeq 6000, and NovaSeq X Plus sequencing and were aggregated using Cell Ranger. Cell doublets were identified and removed using Scrublet (v0.2.3) ^107^ with default parameters except for min_gene_variability_pctl (used 50). Cell barcodes likely containing astrocytes (those with >5% reads aligned to the mouse genome) were also removed. Furthermore, the remaining individual cells were assessed for data quality using gene expression metrics calculated from the raw UMI count matrix. Cells meeting any of the following criteria were flagged as low-quality and removed prior to normalization: (1) total gene expression UMI counts < 2,000, (2) mitochondrial gene reads ≥ 20% of total UMI counts, or (3) fewer than 1,300 genes detected (UMI > 0). These filters retained 21,187 cells for downstream analysis.

#### Feature selection

Normalized gene expression matrices were generated by dividing raw UMI counts by the median total counts per cell and applying log1p transformation. Genes were filtered to retain those expressed in ≥ 3 cells, reducing the feature set to 27,729 genes. Highly variable genes (n = 2,000) were identified and selected for downstream dimensionality reduction, using batch-corrected variance calculations stratified by sequencing library.

### Dimensionality Reduction and Clustering

Principal component analysis (PCA) was applied to the 2,000 highly variable genes, and the first 50 principal components were retained. K-nearest neighbor graphs (k = 15, default) were constructed in PCA space. Cell clustering was performed using the Leiden community detection algorithm (Scanpy implementation; resolution = 0.8, n_iterations = −1, random_seed = 0) to identify transcriptionally distinct cell populations. PAGA (Partition-based graph abstraction) was applied to infer differentiation trajectories and summarize relationships among clusters. UMAP dimensionality reduction was subsequently computed, initializing from PAGA coordinates to preserve trajectory information in visualization.

### Differential expression analysis between mono- and co-cultured neurons

Differential gene expression analysis between mono- and co-cultured neurons was performed using Scanpy’s implementation of the Wilcoxon rank-sum test (Mann-Whitney U test). The analysis was implemented via sc.tl.rank_genes_groups() applied to log-normalized gene expression counts. Genes with adjusted p-value (padj) < 0.01 (n=6921) using the default correction method were retained for comparison with the bulk RNA-seq results.

### Pseudobulking and correlation with a human brain cell atlas

scRNA-seq cell-by-gene count data of excitatory neurons from the publication ^34^ was downloaded from the UCSC Cell Browser. The data was pseudobulked by grouping cells according to age range of the donor, then randomly splitting cells within each group into three technical replicates and computing the mean gene expression across cells within each replicate. Similarly, the scRNAseq data of mono- and co-cultured neurons we generated were pseudobulked by computing the mean raw UMI counts across all cells within each culture condition. Pseudobulk gene expression profiles from the experimental dataset and published developmental brain dataset (Velmeshev et al.) were first subset to 17,092 common genes, normalized to transcripts per million (TPM), and then pairwise Pearson correlation coefficients were computed between experimental samples (culture conditions) and reference samples (grouped by age range of the donor).

### Inference of ligand-receptor pairs between astrocytes and excitatory neurons from single-cell brain atlas using CellChat

Processed and annotated data from a recently published cortical development single-cell RNA-seq study was downloaded^34^. Only cells with prenatal “Age_Range” values were kept. The filtered annData object was further processed with CellChat (v2.2.0), using the latest CellChatDB v2 available for humans. In order to run CellChat, gene symbols in the human CellChatDB were mapped to ENSEMBL IDs using the gprofiler2 (v0.2.3) R package. Then, the recommended CellChat workflow was applied, grouping the cells by the annotated “Lineage”. Briefly, the program identifies over expressed genes and interactions between cell groups, computes the probability of communication at the cell and signaling pathway levels, and computes an aggregated network based on those. Finally, ligand/receptor genes in significantly enriched (p-value <0.1) signaling pathways between astrocytes and excitatory neurons were selected.

### Perturb-seq analysis

#### Data preprocessing and quality filtering of cells

A pipeline similar to that described in Methods section ‘scRNA-seq analysis of mono- and co-cultured neurons’ was used to preprocess data using Cell Ranger and to remove low-quality cells based on gene expression data metrics. Importantly, the specific thresholds were determined on a per-dataset basis. In addition to the pipeline described before, an additional step of single-cell sgRNA assignment was performed with CLEANSER (v1.2) ^47^. Cells with 0 sgRNA assignment were removed from downstream analysis. Notably, predicted doublets were not removed from downstream analysis, as doublets containing a perturbation still contain the transcriptional response elicited by that perturbation. Moreover, for the CRISPRa promoter screen, we also down-sampled the cells containing a MYC sgRNA to match the mean of the rest of the perturbations since MYC activation resulted in a ∼10x increase in cell number that could result in strong bias in downstream transcriptional response analyses. For the CRISPRa pRE screen, we also down-sampled cells containing MYOD1 DDR- or CXCR4-promoter targeting sgRNAs, both positive controls that are over-represented, to match the cell coverage of the rest of the perturbations. Furthermore, for the same screen, we observed and removed expanded single-cell clones with >25 cells as previously described ^108^. After the above filters, 103476, 90577, 67176, and 70459 single cells were retained for the CRISPRi pRE, CRISPRa pRE, CRISPRi promoter, and CRISPRa promoter screens, respectively.

### Building perturbation and random sampling gene expression distributions from Perturb-seq datasets with pySpade

We used pySpade (v0.1.7) ^46^ to analyze the transcriptional response to perturbation in the Pertrurb-seq experiments. Before downstream statistical tests were run, individual genes’ expression distributions were built for each perturbation as well as random samplings of cells in the dataset. For perturbations, the command pySpade DEobs was run, grouping all cells with any sgRNA for a specific target (with the ‘cpm’ normalization method; with cells containing a non-targeting sgRNA as background for the promoter Perturb-seq experiments, or with ‘all other cells’ as background for the pRE Perturb-seq experiments due to high MOI). For random samplings, 1000 samplings were performed by running pySpade DErand (with the ‘sgrna’ randomization method and matched background selection method as DEobs) for the following five cell numbers: 50%, 75%, 100%, 125%, and 150% of the median cell coverage of all perturbations in the same experiment. In downstream analysis (pySpade local and pySpade global), the perturbations were automatically compared to the random sampling cell number that they are closest to.

### Analysis of local transcriptional response to pRE perturbations

For the local transcriptional response to the pRE perturbations where the perturbation-response distance is < 1Mb, the command pySpade local was run on the perturbation and random sampling distributions for the pRE Perturb-seq datasets. FDR correction was applied independently within each target region using the Benjamini-Hochberg method (α = 0.01). Fold change values were filtered to retain only transcriptional responses of >10%. Gene’s CPM was filtered to be > 1 without perturbation (for the CRISPRi dataset) or with perturbation (for the CRISPRa dataset) to remove artificially inflated effect sizes due to low expression. Perturbation-response links that passed these filters were considered significant links.

### Analysis of global transcriptional response to pRE and promoter perturbations

For the global transcriptional response to the pRE and promoter perturbations, the command pySpade global was run on the perturbation and random sampling distributions for all Perturb-seq datasets. FDR correction was applied independently to significance scores within each target region using the Benjamini-Hochberg method (α = 0.01). No fold change filtering was applied to global results. Gene’s CPM expression without perturbation was filtered to be > 1 for all datasets to remove artificially inflated effect sizes due to low expression. For downstream analysis, we focused on CRISPRi/a promoter perturbations that significantly down- or up-regulated the intended target gene, respectively, and fRE perturbations (i.e., pREs linked to a local target gene). We further filtered to only include those that have more than 3 global responsive genes (i.e., DEGs). By these criteria, 88 perturbations across the 4 screens were included, including 10 positive controls.

### Pairwise Spearman correlation of global transcriptional responses

Focusing on the perturbations of interest (detailed in **Methods** ‘Analysis of global transcriptional response to pRE and promoter perturbations), we performed pairwise Spearman correlation on the log2(fold change) values of these perturbations on 247 genes that are found to be responsive to at least one perturbation across at least 3 of the 4 Perturb-seq datasets. The purpose of this filter was to select for genes that are less likely to be driven by batch effect.

### Computation of in vivo mimicry score of CRISPRi/a perturbations

Pseudobulk CPM profiles for 88 perturbations from pySpade global and matched random sampling negative controls were compared to Velmeshev et al. ^34^ excitatory neuron pseudobulks (detailed in **Methods** section ‘Pseudobulking and correlation with a human brain cell atlas’) by first intersecting genes (12,336 in common) and converting the Velmeshev et al. profiles to CPM (column-wise scaling to 1e6). Pairwise Pearson correlations were computed between each perturbation/negative control profile and each Velmeshev et al. reference across the age range of donor. To isolate perturbation signals, correlations were background-normalized per screen by subtracting the mean correlation of negative control profiles, and an average score across age ranges of donor was computed.

### Gene Ontology over-representation analysis perturbations of interest

GO Biological Process over-representation analysis for each perturbation was run by taking that perturbation’s DEG set from the pySpade global results (using the fold-change screen cutoffs of fold change < 0.9 or > 1.1), and testing it against a custom background comprising the genes detected across all perturbations. Enrichment was performed with gseapy’s implementation of Enrichr ^101^ using the human GO_Biological_Process_2025 library (falling back to GO_Biological_Process_2023 when needed), Enrichr’s Benjamini–Hochberg FDR adjustment, and a significance threshold of adjusted P < 0.01, with results saved as standard Enrichr report files per perturbation.

### Disease gene set over-representation analysis for bulk astrocyte-responsive neuron gene modules and downstream genes of CRISPRi/a perturbations

22 total publications containing gene sets relevant to central nervous system-related disorders were selected (for a full list of references, see **Table S16**). Over-representation of the disease gene sets in astrocyte-responsive neuron gene modules was analyzed using a three-stage preprocessing and testing workflow: In stage 1, a ‘gene universe’ was prepared by identifying protein-coding genes expressed in our bulk RNA-seq dataset (baseMean>=3; n=15332 for neuron genes, n=13565 for astrocyte genes). In stage 2, disease gene sets from publications were extracted genes meeting publication-specific significance thresholds (typically FDR- or adjusted p < 0.01–0.05, detailed in **Table S17**) and stratified into up- and down-regulated for case/control gene expression studies. Only disease genes that intersect with the gene universe were retained. In stage 3, gseapy’s (v1.1.9) implementation of Enrichr was applied to each gene module using the above disease gene sets, with the background as the ‘gene universe’ (protein-coding genes meeting expression filters and inclusion criteria), Benjamini–Hochberg FDR correction, and significance threshold of adjusted p < 0.01. The same analysis was also run on the downstream gene sets of the CRISPRi/a perturbations of interest.

### RT-qPCR analysis

Three biological replicates per sgRNA (each sgRNA-biological replicate combination is termed a biological sample) and two technical replicates per biological sample were performed. Within each technical replicate, the difference in Ct values of the target gene and the housekeeping control gene TBP was calculated and the relative expression of the target gene was calculated as 2*Δ*^Ct^ (*Δ*Ct = Ct(TBP) - Ct(target gene). The mean across technical replicates was computed for each biological sample for statistical analysis. The 2*^Δ^*^Ct^ values were compared between a targeting sgRNA and the non-targeting sgRNA using unpaired t tests. P values < 0.1 were considered significant.

### MEA analysis

Spike files generated by the AxIS Navigator software were imported into AxIS Neural Metric Tool (v4.1.1) to generate raster plots for individual wells, and to compute and export recommended neural metrics for further analysis. For each metric, Mann-Whitney U tests were performed at each time point comparing each perturbation condition (sgRNAs, CRISPRa/i) against the sgNT2 control (**Table S17**). Multiple testing correction was applied across all comparisons using Bonferroni method, with significance thresholds at p < 0.05, p < 0.01, and p < 0.001; results were visualized as heatmaps displaying differences from control (median values subtracted), with significance stars annotating Bonferroni-corrected p-values, rows ordered by experimental condition.

### Immunofluorescence image processing

Raw image files were processed with Fiji^109^. CZI files were batch-imported with Bio-Formats in Hyperstack mode (XYCZT stack order)^110^, and for z-stacked images, maximum intensity projections were computed per channel, whereas single-slice images were processed directly without projection. Each channel was isolated and assigned a pseudocolor, and display intensity ranges were normalized using fixed maximum values (typically 65,535 for 16-bit depth, with channel-specific adjustments for optimal visibility across experiments). A scale bar (100 µm width) was added to all outputs. Images were converted to RGB to preserve pseudocolors, and composite images were generated by merging channels for multi-marker visualization. Neurite outgrowth was quantified by segmenting MAP2-stained images using a fixed intensity threshold (5000) following rolling-ball background subtraction, while nuclei were identified from DAPI channels using Otsu thresholding and watershed separation. The total neurite surface area was measured as a percentage of the field of view from the resulting binary masks. Finally, this area fraction was normalized to the total nuclei count per image to derive the average neurite area per cell. The effects of perturbations were compared to sgNT2 using one-way ANOVA following Dunnett’s multiple comparisons test. Adjusted p values were reported.

### Use of LLM in the writing of this manuscript

The writing of some parts of **Methods** was assisted by LLM through summarizing existing protocols, lab notes, and code. Every result was human-edited and -verified.

## Notes

https://data.igvf.org

